# Studying evolutionary dynamics across the entire Sequence Read Archive

**DOI:** 10.64898/2026.11.23.768713

**Authors:** Andrew C. Seacord, Rayan Chikhi, Danielle Callan, Steven Weaver, Patrik Smeds, Sergei L. Kosakovsky Pond, Anton Nekrutenko

## Abstract

Querying the Sequence Read Archive traditionally relied on submitter metadata that is often missing, inconsistent, or wrong. Logan changed that picture by assembling the archive for direct sequence searches with kmindex and LexicMap. Yet hits are not alignments. Variation and uneven coverage fragment genes across assembly pieces, while redundant outbreak isolates overwhelm comparative evolutionary tools. We built disassembler to reconstruct complete coding sequences from unitig graphs or contigs, thin clonal duplicates, and build deep cohorts spanning up to millions of runs. From 4.99 million SARS-CoV-2 hits, disassembler recovered 910,848 strict and 1,153,319 permissive Spike sequences. Paired with the Aeon tools (HyphAeon, RhizAeon, and ChronAeon), downsampled cohorts (≤ 200 haplotypes) resolve evolutionary dynamics in seconds without iterative tree search. Across SARS-CoV-2, microbiome obelisks, and *Helicobacter pylori*, panels recover a 2.55-fold Omicron clock acceleration, *gyrA* drug-pocket rigidity, *oipA*/*vacA*/Spike co-selection, 57.4% mixed obelisk infections with six mosaic tracts, and 5.37-fold *vacA* recombination enrichment.

## Introduction

Until recently, biologists could not routinely mine the full Sequence Read Archive (SRA) for arbitrary genes and uncatalogued variants. Built as an international public repository of raw sequencing reads (Kodama et al., 2012), the archive now exceeds 47 million experiments (Chikhi et al., 2025), yet finding runs of interest long meant querying submitter metadata such as organism names, hosts, tissue types, and collection dates (Gonçalves and Musen, 2019). That metadata is notoriously unreliable. Required fields are routinely blank, sample descriptions use inconsistent terms, and contaminated or mislabeled runs carry the wrong taxonomic names (Gonçalves and Musen, 2019; Huang et al., 2025). Metadata gaps left much of the SRA dark to systematic comparative genomics.

Petabase-scale sequence indexes began opening that archive before full read assembly became practical. Serratus enabled cloud-scale viral RNA-dependent RNA polymerase screening (Edgar et al., 2022), while containment sketch search in sourmash and branchwater (Irber et al., 2022), full-text *k*-mer graph indexing in MetaGraph (Karasikov et al., 2025), universal seed indexing in Pebblescout (Shiryev and Agarwala, 2024), and alignment-free phylogenetic placement in UShER (Turakhia et al., 2021) alongside sketching workflows (Faure et al., 2026) demonstrated rapid discovery across massive public collections. The Logan project extended this shift to near-complete archive assembly, collapsing each sequencing run into unitigs and contigs (see Methods) (Chikhi et al., 2025). This representation shrinks raw data roughly tenfold into unitigs that preserve intra-sample genetic variation and roughly fiftyfold into contigs that trade minor alleles for longer sequences. Two indexing engines on Galaxy (The Galaxy Community, 2024), kmindex (Lemane et al., 2022, 2024) over Bloom-filter unitig matrices and LexicMap (Shen et al., 2026) over lexicographic contig seeds, let any biologist search these assemblies by sequence alone at https://usegalaxy.org.

Search hits alone still stop short of evolutionary inference, because natural variation and uneven read depth fragment genes across multiple assembly pieces (Compeau et al., 2011; Nurk et al., 2017). Neither read-free *k*-mer containment tables nor local pairwise contig hits yield the clean, frame-verified multiple sequence alignments needed downstream. We built disassembler to bridge that gap. Given a gene or region of interest, disassembler consumes raw kmindex or LexicMap outputs, reconstructs full coding sequences from unitig subgraphs or tiled contigs, filters frameshifts and paralogous mixtures, thins near-identical duplicates, and emits alignments ready for comparative analysis.

This cohort builder pairs naturally with a new family of fast evolutionary tools—HyphAeon (Kosakovsky Pond et al., 2026c), RhizAeon (Kosakovsky Pond et al., 2026a), and ChronAeon (Kosakovsky Pond et al., 2026b), described in companion method preprints and referred to collectively here as the Aeon suite. Classical selection, recombination, and molecular-clock frameworks—PAML (Yang, 2007), HyPhy methods including MEME, BUSTED, and GARD (Kosakovsky Pond et al., 2005; Murrell et al., 2012, 2015; Kosakovsky Pond et al., 2006), and BEAST (Suchard et al., 2018; Bouckaert et al., 2019), alongside fast post-tree dating tools such as TreeTime (Sagulenko et al., 2018) and LSD2 (To et al., 2016)—either require iterative phylogenetic search or depend on an accurate precomputed tree (Morel et al., 2021; Turakhia et al., 2021). The Aeon tools avoid tree search: HyphAeon builds an internal neighbor-joining tree when no external topology is supplied, whereas RhizAeon and ChronAeon compute directly from aligned sequence patterns and pairwise distances. Here we benchmark disassembler cohort assembly at archive scale up to 4.99 million runs and evaluate downstream Aeon analyses on downsampled panels of up to 200 sequences across pandemic SARS-CoV-2, uncultured microbiome obelisks, and *Helicobacter pylori*.

## Results and Discussion

### System Architecture

Our workflow begins on public Galaxy servers (The Galaxy Community, 2024), where two complementary search engines query the Logan assemblies without touching submitter metadata (Figure 1A). The first engine, kmindex, indexes Logan unitigs as inverted Bloom filter matrices (Bloom, 1970) built with kmtricks (Lemane et al., 2022, 2024). A Bloom filter is a compact bit array that tests whether a sequence word occurs in a dataset without storing the sequence itself. Stacking one filter per SRA run into an inverted matrix lets kmindex test millions of runs at once, while the findere overlapping-word scheme suppresses false hits (Robidou and Peterlongo, 2021) (see Methods). Because tens of millions of runs cannot fit inside one monolithic file, kmindex splits the archive by sequencing assay and broad organism group—such as genomic, metagenomic, transcriptomic, and viral datasets—and distributes each group across smaller index partitions called shards. That sharded layout lets Galaxy scan the entire SRA in parallel or target specific slices such as viral or human microbiome shards directly. A biologist submits a query sequence—for example, a viral gene or a bacterial locus—and kmindex breaks that query into short overlapping sequence words (*k*-mers) and checks which SRA runs contain them. The tool returns a table listing each matching SRA accession alongside its *k*-mer match score, reporting the fraction of query words present in that run.

**Figure 1:**
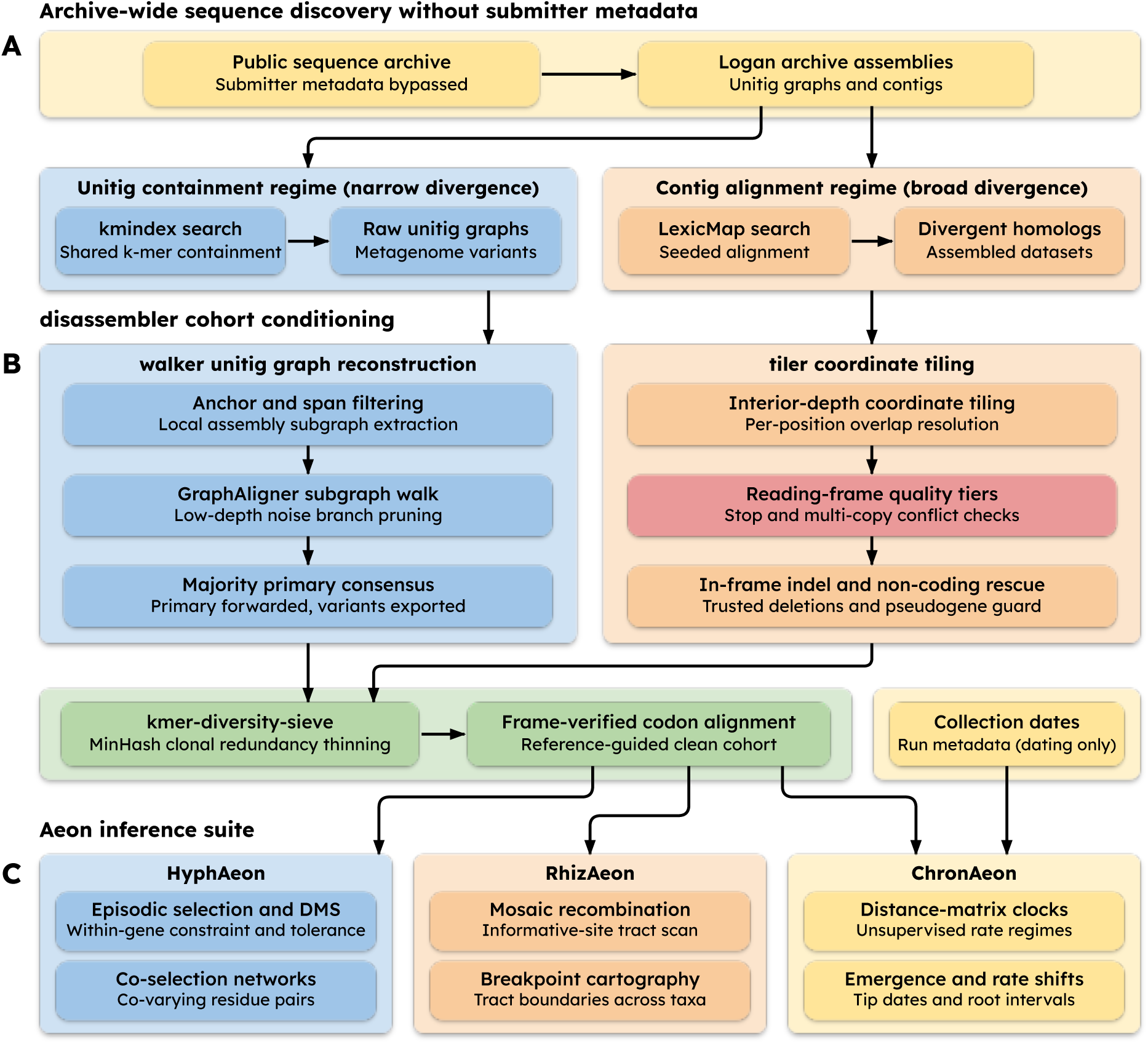
Overview of the workflow connecting archive-wide sequence search, disassembler cohort assembly, and the Aeon evolutionary suite. (A) Logan assemblies feed two search routes on public Galaxy servers: kmindex exact short-word matching over unitig graphs for close relatives and metagenomes, and LexicMap seeded alignment over contigs for more divergent homologs. (B) disassembler turns raw hits into alignments through walker (local graph traversal, error pruning, and majority sequence assembly with minor variants saved separately) or tiler (coordinate stitching, reading-frame and conflict checks, and indel rescue), followed by MinHash redundancy thinning (kmer-diversity-sieve) and codon alignment. (C) Clean alignments and collection dates feed HyphAeon (within-gene constraint, selection, and co-evolving sites), RhizAeon (recombination tracts), and ChronAeon (molecular clocks and rate regimes). This schematic uses a muted pastel palette distinct from the Okabe-Ito data palette in Figures 2 to 4: cream marks archive sources and clocks, sky blue marks the unitig route and HyphAeon, peach marks the contig route and RhizAeon, coral marks reading-frame QC, sage marks thinning and alignment, and black arrows show data flow. Abbreviations: QC, quality control.

The second engine, LexicMap, organizes Logan contigs into a hierarchical lexicographic seed index partitioned across broad organism and assay categories for direct sequence alignment (Shen et al., 2026). Instead of requiring exact full-length word matches, LexicMap indexes sorted variable-length prefix and suffix seeds so mutated loci still anchor alignment chains (see Methods). When a biologist submits the same query sequence, LexicMap searches against millions of assembled SRA datasets and returns a BLAST-like alignment table. Each row gives the matching SRA accession, the aligned coordinates on both the query and the target contig, percent sequence identity, and the pairwise alignment.

The two engines answer different biological questions and cover different ranges of sequence divergence. In practice, kmindex gives a fast presence-or-absence answer: it tells a biologist which SRA runs carry sequence words from a query, without returning an alignment. Because exact short-word matching breaks down as mutations accumulate, kmindex works best when searching for close relatives at low sequence divergence, or when polymorphic raw-read graphs fail to assemble into clean contigs. LexicMap, by contrast, returns explicit local alignments and tolerates many more nucleotide substitutions. As a result, LexicMap finds much more divergent homologs across assembled datasets.

Downstream evolutionary tools do not take search hits. They take alignments, so we built disassembler to turn raw kmindex and LexicMap outputs into clean multiple sequence alignments (Figure 1B). Mirroring the two search indices, disassembler has two assembly paths: walker for kmindex results and tiler for LexicMap results.

When starting from kmindex, a biologist has a list of matching SRA accessions without assembled gene sequences. The walker component (implemented as logan-walker) streams each matching run’s compressed unitig graph directly from Logan, extracts the local subgraph matching the query, and traces a continuous path through it (Rautiainen and Marschall, 2020). Because graph aligners otherwise tend to pick rare sequencing-error branches that happen to match the reference sequence, walker trims away very low-depth branches and selects the majority allele at polymorphic forks. It outputs one clean primary consensus sequence per matching SRA accession while writing minor within-sample variants to separate files so intra-host diversity is preserved without distorting cross-sample alignments.

When starting from LexicMap, local alignments already exist, yet a single gene is often split across multiple contigs or interrupted by deletions, frameshifts, or multi-copy collisions. The tiler component reads the alignment table accession by accession and tiles all matching pieces onto the query coordinate frame. Where pieces overlap, it favors bases near the interior of an alignment over noisy ends. It checks gene coverage, recognizes genuine in-frame deletions without dropping them as missing data, flags frameshifts, premature stop codons, and conflicting multi-copy overlaps, and writes the clean sequences into a multiple sequence alignment.

Archive-wide searches often return hundreds of thousands of near-identical isolates from dense outbreak sampling, which inflate runtime without adding new evolutionary information (Morel et al., 2021). To remove that redundancy, disassembler includes a diversity sieve (wrapping kmer-diversity-sieve). The sieve scans sequences in order and keeps a sequence only when it differs sufficiently from every representative already retained, dropping clonal duplicates while keeping rare variants intact.

Finally, the alignment step locks clean coding sequences into frame-verified codon alignments, reverse-complementing opposite-strand hits and filtering short overhanging fragments (see Methods). For non-coding queries, codon checks turn off automatically so the pipeline still produces clean nucleotide alignments.

Those conditioned alignments then feed straight into the Aeon suite (Figure 1C). HyphAeon takes a codon alignment and uses a trained neural model to report episodic selection, within-gene residue plasticity, and co-evolving codon pairs in seconds, deriving a neighbor-joining tree internally when no topology is given (Kosakovsky Pond et al., 2026c). RhizAeon detects where recombination has swapped sequence segments between lineages directly from aligned site patterns (Kosakovsky Pond et al., 2026a). ChronAeon pairs aligned sequences with collection dates to estimate substitution rates, emergence dates, and rate regimes from pairwise distances without Bayesian tree sampling (Kosakovsky Pond et al., 2026b). All Aeon case studies below ran on disassembler v0.1.0 cohorts and were audited for haplotype concordance against final v0.2.0 builds without rerunning the Aeon tools on v0.2.0 outputs. All parameters and provenance appear in Methods.

### SARS-CoV-2

To test cohort assembly at pandemic scale, we queried the SRA for the surface spike glycoprotein gene of severe acute respiratory syndrome coronavirus 2 (SARS-CoV-2), which engages the human ACE2 receptor and drives neutralizing antibody escape (Lan et al., 2020). Table 1 and Figure 2 summarize every stage of this five-million-run pipeline.

**Figure 2:**
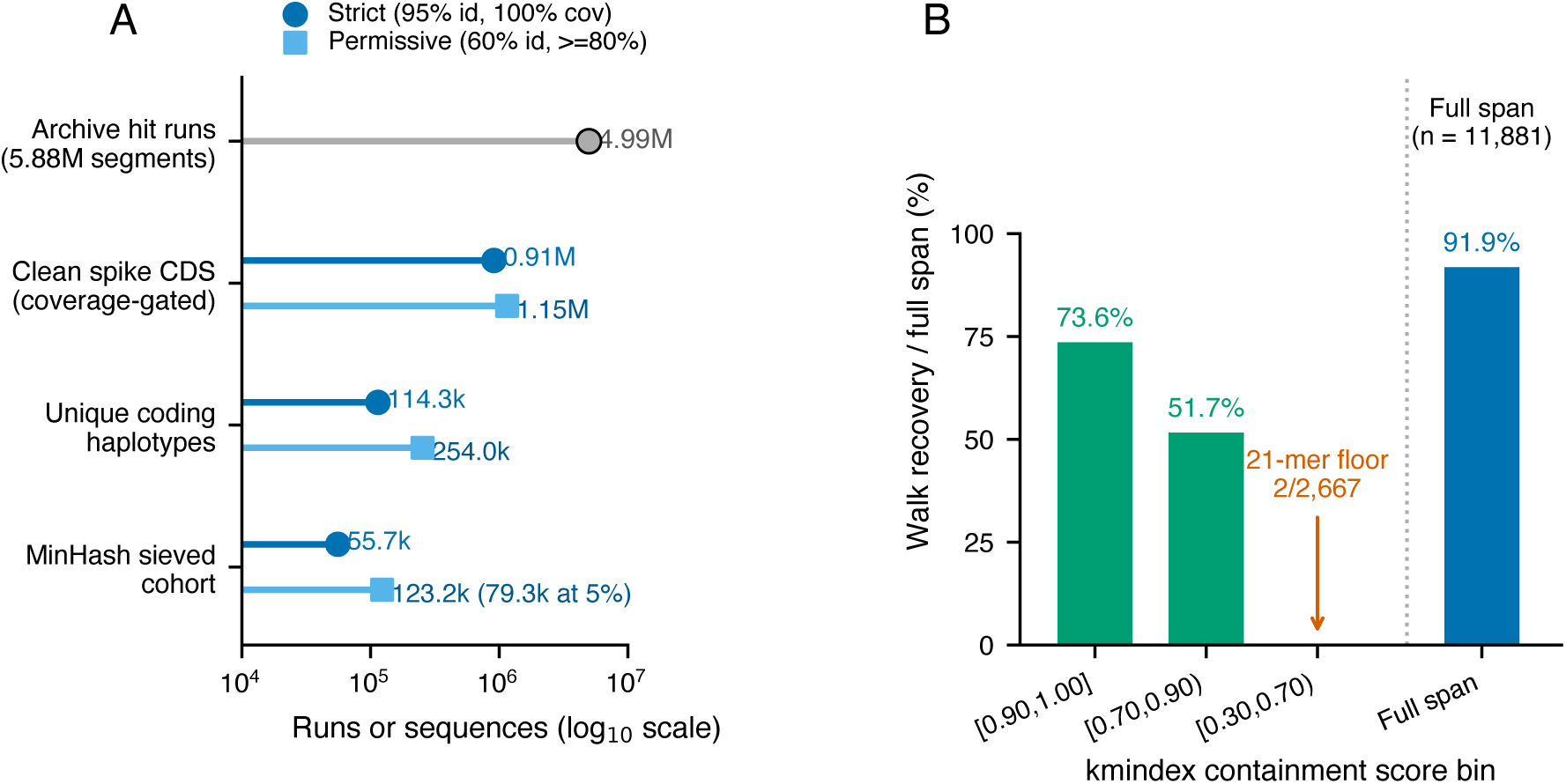
Archive cohort recovery and conditioning across both search routes. (A) SARS-CoV-2 spike LexicMap contig funnel across 4,987,809 hit runs (5,878,944 segments) on a log_10_ scale. Strict tiling yields 910,848 clean sequences, 114,292 haplotypes, and 55,723 sieved representatives (Distinct excludes and Sieved includes the kept reference). Permissive tiling yields 1,153,319 clean sequences, 253,987 haplotypes, and 123,176 sieved representatives under a 25% gap cap (79,255 under 5%). (B) Stratified kmindex-to-walker recovery across 20,000 sampled viral runs grouped into high, medium, and low containment bins against the ancestral spike query, showing the drop-off in the lowest match tier (2/2,667) alongside 91.9% full-span recovery (*n* = 11, 881). Black-rimmed gray marks archive hits, dark blue marks strict cohorts and full span, sky blue marks permissive cohorts, bluish green marks ancestral walks, and vermilion marks the low-containment floor. Abbreviations: CDS, coding sequence.

**Table 1:** Archive-wide cohort discovery, reconstruction, and compression across three systems with disassembler v0.2.0. In the tiler route, assembled-contig cohorts stream LexicMap alignment tables directly into codon coordinates, rescuing in-frame deletions and stitched multi-segment alignments while separating frameshifts, premature stop codons, and conflicting segment overlaps across millions of assemblies. In the walker route, raw-unitig cohorts rank sequencing runs by kmindex containment and thread local de Bruijn assembly subgraphs (pinned GraphAligner 1.0.17b) while recording intra-sample alternative haplotypes. Canonical 15-mer MinHash sketching (*τ* = 0.005, sketch size 256) compresses clonal redundancy. Two-part wall times (*A* + *B* min) report tiler cohort conditioning (*A*) plus MinHash sieving (*B*), whereas single values report total cohort conditioning or end-to-end reconstruction time. Superscript a denotes Distinct excluding and Sieved including the kept ancestral reference, b excludes the reference from both, c marks shared-node execution alongside concurrent 48-worker unitig walks, d notes historical v0.1.0 comparators (76,360 strict SARS-CoV-2 Spike sequences and 34/345 clipped obelisks), and e reports an alternate 5% gap-cap permissive sieve.

| Biological target | Search and assembly route | Candidate runs | Clean cohorts | Distinct / sieved | Wall time | Key quality-control and biological outcome (v0.2.0) |
|---|---|---|---|---|---|---|
| SARS-CoV-2 Spike (3,822 bp) | LexicMap + tiler (strict) | 4,987,809 | 910,848 | 114,292 / 55,723<sup>a</sup> | 78 + 5.5 min | Recovers 836,270 indel-bearing runs<sup>d</sup>, flags 1,657 frameshifts, 2,537 stops, and 59 mixed overlaps |
| SARS-CoV-2 Spike (3,822 bp) | LexicMap + tiler (permissive) | 4,987,809 | 1,153,319 | 253,987 / 123,176<sup>a</sup> | 282<sup>c</sup> + 16.5 min | Rescues 32,319 split multi-segment runs plus 538,996 clean partials (79,255 at 5% gap cap<sup>e</sup>), flags 164 mixed overlaps |
| SARS-CoV-2 Spike (ancestral reference) | kmindex + walker (20k walked) | 5,138,371 (20,000 walked) | 11,881 (10,914 full span) | 7,049 / 5,178<sup>b</sup> | 158 min | 51.7% recovery in [0.70, 0.90) containment bin as divergent Omicron runs retain conserved 21-mer anchors, 20,390 alternatives |
| *Helicobacter pylori* (25-locus panel) | LexicMap + tiler (20 strict, 5 loose) | 2,620–26,260 per locus | 2–2,466 per locus | 2–903 / not sieved | 36 s | *fliD* restores 2,040 pseudogene-anchored rows, *gyrA* retains 1,605 via 3-bp deletion rescue, loose adhesins flag 4,306 paralog mixes |
| Microbiome obelisks (5 lineages) | kmindex (141 shards) + walker | 924 | 350 | 311 / not sieved (+931 alternatives) | 27.4 min | Zero 3′ terminal clipping (0/350<sup>d</sup>), flags 57.4% (201/350) intra-sample mixtures, drops 16 overhang rows in antisense *oblin2* |

On the LexicMap side, searching assembled contigs returned nearly five million matching SRA runs (4,987,809 runs across 5,878,944 alignment segments). Passing that table through tiler under strict full-length criteria recovered 910,848 clean spike coding sequences in 78 minutes (Figure 2A). More than 91% of those clean pandemic spike sequences (836,270 runs) carry real in-frame deletions in the N-terminal domain or Omicron sublineages, and tiler counts internal deletions as valid sequence span instead of discarding deletion-bearing lineages as incomplete. Meanwhile, tiler isolated frameshifts, premature stop codons, and conflicting mixed overlaps into separate audit files. Collapsing identical sequences reduced those 910,848 clean runs to 114,292 distinct spike haplotypes, and running the diversity sieve compressed them further to 55,723 representatives in 5.5 minutes—nearly a ninetyfold reduction from the initial archive hits. Under permissive settings allowing partial assemblies, tiler recovered 1,153,319 clean sequences (253,987 distinct haplotypes, thinned to 123,176 representatives under a 25% gap cap or 79,255 under 5%) and stitched together 32,319 spike genes split across multiple contigs.

On the kmindex side, querying viral unitig shards with the ancestral spike sequence returned 5,138,371 candidate runs. To measure how graph walking behaves across increasing divergence, we drew a stratified sample of 20,000 runs across high, medium, and low *k*-mer containment bins and ran walker (Figure 2B). Overall, walker reconstructed 11,881 spike sequences (91.9% spanning the entire gene) plus 20,390 minor intra-sample variant paths, and the sieve compressed 7,049 unique primary sequences down to 5,178 representatives in 25 seconds. Recovery reached 73.6% in the highest match tier and remained at 51.7% in the [0.70, 0.90) containment tier, where divergent Omicron isolates still shared enough intact 21-mer anchors with the ancestral spike query to walk cleanly. In the [0.30, 0.70) tier, however, recovery dropped to 0.07% (2 of 2,667 runs). Both high divergence and fragmented low-depth viral sequencing in mixed samples break shared 21-mer spans, marking the practical boundary where biologists switch to LexicMap.

Downstream Aeon benchmarks were performed on downsampled cohorts of at most 200 taxa per analysis rather than the full sieved panels (Table 2). For spike, we analyzed 161 aligned sequences: the top 160 global LexicMap haplotypes representing 545,898 v0.1.0 permissive runs across 15 countries plus the ancestral reference (140 of 160 carry in-frame deletions). In 29.5 seconds, HyphAeon detected 27 of 110 variable codons at nominal *p* ≤ 0.05 against 5.5 expected by chance, alongside 171 of 5,995 variable pairs at *q* ≤ 0.05 across seven network sectors (Figure 3B, Table 2). None of the 27 codons survived Benjamini–Hochberg correction across all 1,274 spike sites, while one codon—multi-allelic receptor-binding escape hub Glu484 (LRT = 16.32, *p* = 1.21 × 10^−4^, carrying Omicron E484A as well as E484K (Starr et al., 2020; Greaney et al., 2021; Harvey et al., 2021))—remained significant under both Bonferroni and Benjamini–Hochberg adjustment restricted to the 110 variable codons (*q*_var_ = 0.0133). The remaining top nominal signals ranked in likelihood-ratio order are class-3 antibody-escape residue Val445 (V445P/A) (Cao et al., 2023), S2 residue Asp1118 (D1118H), fusion-subunit site Asp950 (D950N/H), signal-peptide residue Ser13 (S13I), and furin-cleavage site Pro681 (P681R/H) (Harvey et al., 2021).

**Figure 3:**
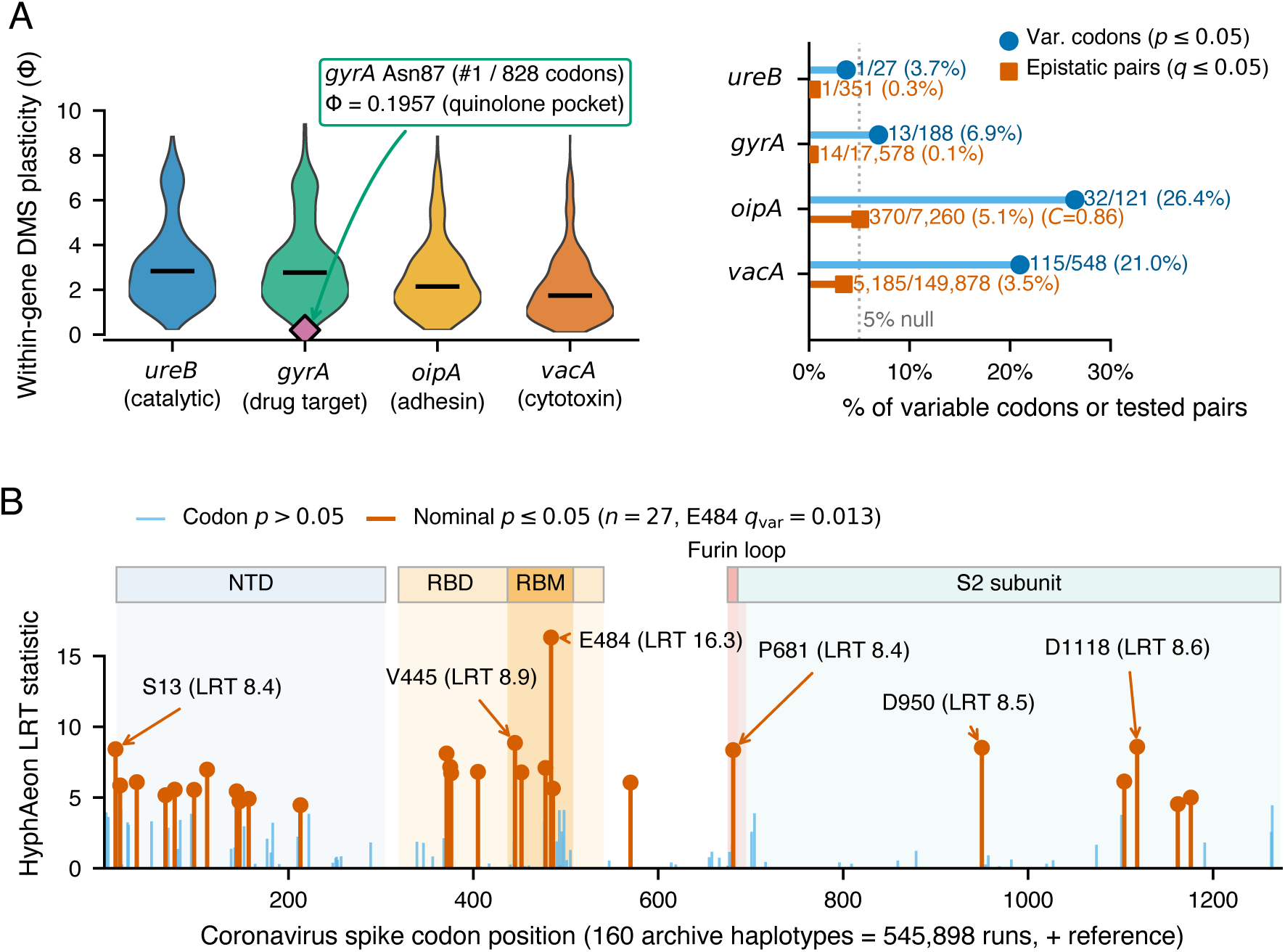
Selection, within-gene mutational plasticity, and co-evolving sites inferred with HyphAeon v0.2.0 on disassembler v0.1.0 downsampled cohorts. (A) Functional comparison across four *Helicobacter pylori* genes (*n* = 200 haplotypes per locus, *n* = 184 in strict *gyrA*). Left violins show per-site within-gene plasticity (Φ, black medians) across urease *ureB*, gyrase *gyrA* (marking quinolone-pocket residue Asn87 as the single most rigid site across all 828 codons), adhesin *oipA*, and cytotoxin *vacA*. Because Φ normalizes variation relative to each alignment’s diversity, pooled distributions are not compared across genes. Right paired lollipops plot the fraction of variable codons at nominal *p* ≤ 0.05 and tested pairs (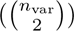) at *q* ≤ 0.05 on a linear scale against the 5% chance expectation (dotted vertical line). (B) Selection scan across the 1,274-codon SARS-CoV-2 spike gene (*n* = 160 global haplotypes representing 545,898 v0.1.0 runs plus reference, 29.5 seconds), aligned under structural domains and labeling top codons E484 (*q*_var_ = 0.0133), V445, D1118, D950, S13, and P681 among 27 nominal sites. Colors follow Okabe-Ito dark blue (*ureB* and variable codons), bluish green (*gyrA*), orange (*oipA*), sky blue (neutral spike codons), vermilion (*vacA*, epistatic pairs, and nominal spike codons), and reddish-purple diamonds (Asn87). Abbreviations: DMS, deep mutational scanning, LRT, likelihood-ratio test, NTD, N-terminal domain, RBD, receptor-binding domain, and RBM, receptor-binding motif.

**Table 2:**
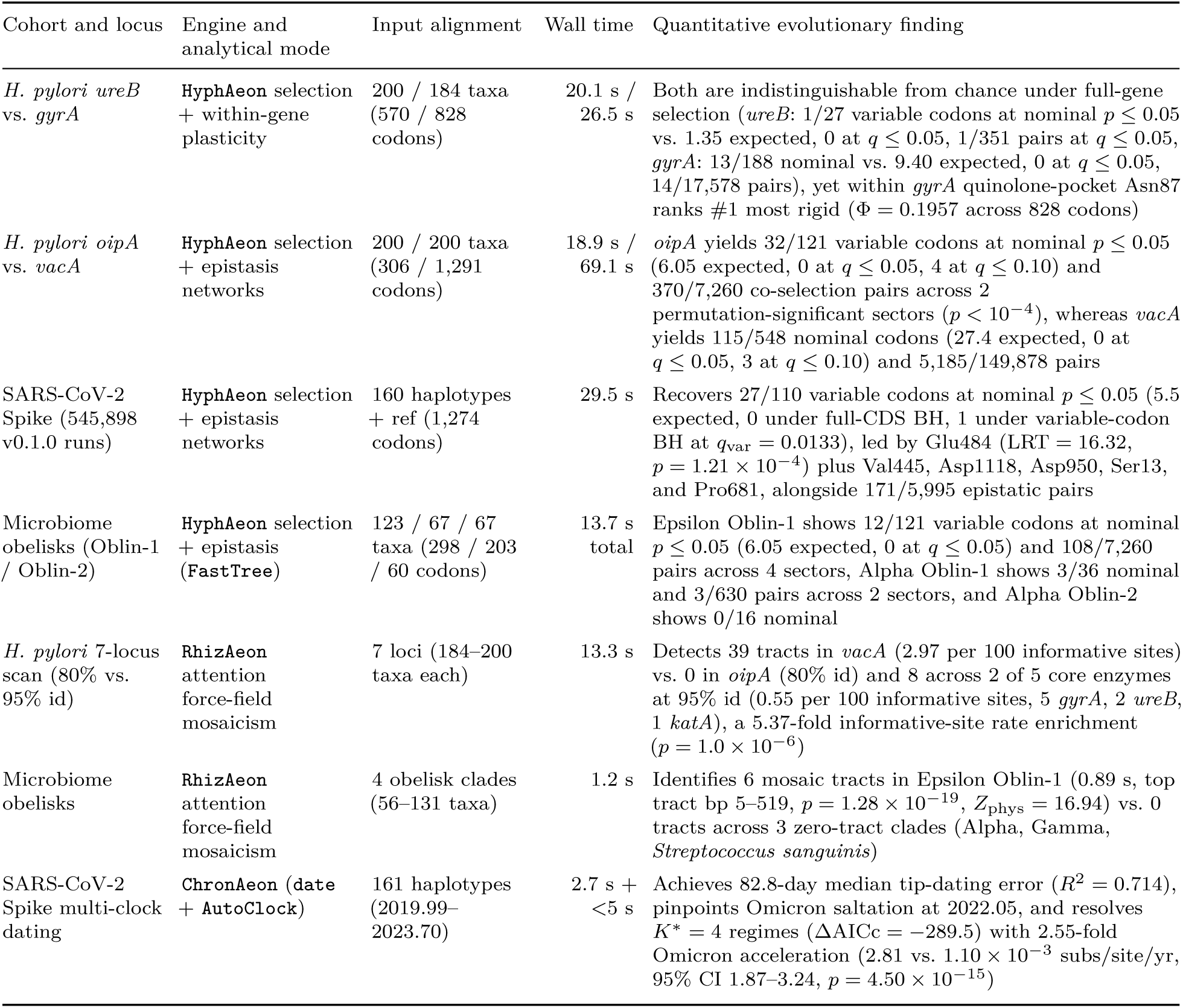
Evolutionary dynamics across downsampled public sequencing cohorts inferred with HyphAeon v0.2.0, RhizAeon v0.1.0b3, and ChronAeon v0.1.0. HyphAeon builds an internal neighbor-joining tree without search for *Helicobacter pylori* and SARS-CoV-2 and accepted FastTree topologies for obelisks, whereas RhizAeon and ChronAeon use sequence patterns and pairwise distances without trees. All input alignments (at most 200 taxa per run) were conditioned with disassembler v0.1.0, and no Aeon analysis has been rerun on v0.2.0 cohorts. Core *H. pylori* cohorts (*ureB*, *gyrA*, *recA*, *atpA*, *katA*) match v0.2.0 clean haplotypes at 99.5% to 100% identity, and 12 of 13 recombinant Epsilon Oblin-1 isolates persist in v0.2.0, though v0.2.0 flags 57.4% (201/350) of obelisk primaries in the full-harvest run as intra-sample mixtures (see Obelisks).

| Cohort and locus | Engine and analytical mode | Input alignment | Wall time | Quantitative evolutionary finding |
|---|---|---|---|---|
| *H. pylori ureB* vs. *gyrA* | HyphAeon selection + within-gene plasticity | 200 / 184 taxa (570 / 828 codons) | 20.1 s / 26.5 s | Both are indistinguishable from chance under full-gene selection (*ureB*: 1/27 variable codons at nominal *p* ≤ 0.05 vs. 1.35 expected, 0 at *q* ≤ 0.05, 1/351 pairs at *q* ≤ 0.05, *gyrA*: 13/188 nominal vs. 9.40 expected, 0 at *q* ≤ 0.05, 14/17,578 pairs), yet within *gyrA* quinolone-pocket Asn87 ranks #1 most rigid (Φ = 0.1957 across 828 codons) |
| *H. pylori oipA* vs. *vacA* | HyphAeon selection + epistasis networks | 200 / 200 taxa (306 / 1,291 codons) | 18.9 s / 69.1 s | *oipA* yields 32/121 variable codons at nominal *p* ≤ 0.05 (6.05 expected, 0 at *q* ≤ 0.05, 4 at *q* ≤ 0.10) and 370/7,260 co-selection pairs across 2 permutation-significant sectors (*p* < 10<sup>−4</sup>), whereas *vacA* yields 115/548 nominal codons (27.4 expected, 0 at *q* ≤ 0.05, 3 at *q* ≤ 0.10) and 5,185/149,878 pairs |
| SARS-CoV-2 Spike (545,898 v0.1.0 runs) | HyphAeon selection + epistasis networks | 160 haplotypes + ref (1,274 codons) | 29.5 s | Recovers 27/110 variable codons at nominal *p* ≤ 0.05 (5.5 expected, 0 under full-CDS BH, 1 under variable-codon BH at *q*<sub>var</sub> = 0.0133), led by Glu484 (LRT = 16.32, *p* = 1.21 × 10<sup>−4</sup>) plus Val445, Asp1118, Asp950, Ser13, and Pro681, alongside 171/5,995 epistatic pairs |
| Microbiome obelisks (Oblin-1 / Oblin-2) | HyphAeon selection + epistasis (FastTree) | 123 / 67 / 67 taxa (298 / 203 / 60 codons) | 13.7 s total | Epsilon Oblin-1 shows 12/121 variable codons at nominal *p* ≤ 0.05 (6.05 expected, 0 at *q* ≤ 0.05) and 108/7,260 pairs across 4 sectors, Alpha Oblin-1 shows 3/36 nominal and 3/630 pairs across 2 sectors, and Alpha Oblin-2 shows 0/16 nominal |
| *H. pylori* 7-locus scan (80% vs. 95% id) | RhizAeon attention force-field mosaicism | 7 loci (184–200 taxa each) | 13.3 s | Detects 39 tracts in *vacA* (2.97 per 100 informative sites) vs. 0 in *oipA* (80% id) and 8 across 2 of 5 core enzymes at 95% id (0.55 per 100 informative sites, 5 *gyrA*, 2 *ureB*, 1 *katA*), a 5.37-fold informative-site rate enrichment (*p* = 1.0 × 10<sup>−6</sup>) |
| Microbiome obelisks | RhizAeon attention force-field mosaicism | 4 obelisk clades (56–131 taxa) | 1.2 s | Identifies 6 mosaic tracts in Epsilon Oblin-1 (0.89 s, top tract bp 5–519, *p* = 1.28 × 10<sup>−19</sup>, *Z*<sub>phys</sub> = 16.94) vs. 0 tracts across 3 zero-tract clades (Alpha, Gamma, *Streptococcus sanguinis*) |
| SARS-CoV-2 Spike multi-clock dating | ChronAeon (date + AutoClock) | 161 haplotypes (2019.99–2023.70) | 2.7 s + <5 s | Achieves 82.8-day median tip-dating error (*R*<sup>2</sup> = 0.714), pinpoints Omicron saltation at 2022.05, and resolves *K*<sup>\*</sup> = 4 regimes (ΔAICc = −289.5) with 2.55-fold Omicron acceleration (2.81 vs. 1.10 × 10<sup>−3</sup> subs/site/yr, 95% CI 1.87–3.24, *p* = 4.50 × 10<sup>−15</sup>) |

Running ChronAeon on those same 161 dated sequences took under 8 seconds (Figure 4C). Without receiving lineage labels, ChronAeon fit an epidemiological crash knot at 2022.05 coinciding with Omicron replacement (Viana et al., 2022) and selected *K*^∗^ = 4 unsupervised clock regimes (ΔAICc = −289.5 over a single clock). Regime 1 (*n* = 79, 0 Omicron haplotypes, spanning 2019.99–2023.04) captured the pre-Omicron ancestral, Alpha, and Delta backbone at *µ* = 1.10 × 10^−3^ substitutions/site/year with an inferred root *t*_MRCA_ of 2019.77 (95% Fieller CI 2019.21–2019.98), consistent with late-2019 pandemic emergence (Pekar et al., 2022). Regime 2 (*n* = 15, 12 Delta plus 2 Alpha and 1 D614G haplotype) resolved an accelerated Delta surge at 2.17 × 10^−3^ substitutions/site/year. Regime 0 (*n* = 58, all 58 carrying Omicron E484A and N501Y) evolved 2.55-fold faster than the backbone (2.81 × 10^−3^ substitutions/site/year, 95% delta-method ratio CI 1.87–3.24, Wald slope difference *p* = 4.50×10^−15^), with a regime root of 2021.37 (CI 2021.19–2021.51) preceding its late-2021 global detection (Viana et al., 2022). Regime 3 (*n* = 9 Delta haplotypes sampled over 11 weeks in 2021) formed a tight clonal cluster with no estimable positive clock (*p* = 0.83).

**Figure 4:**
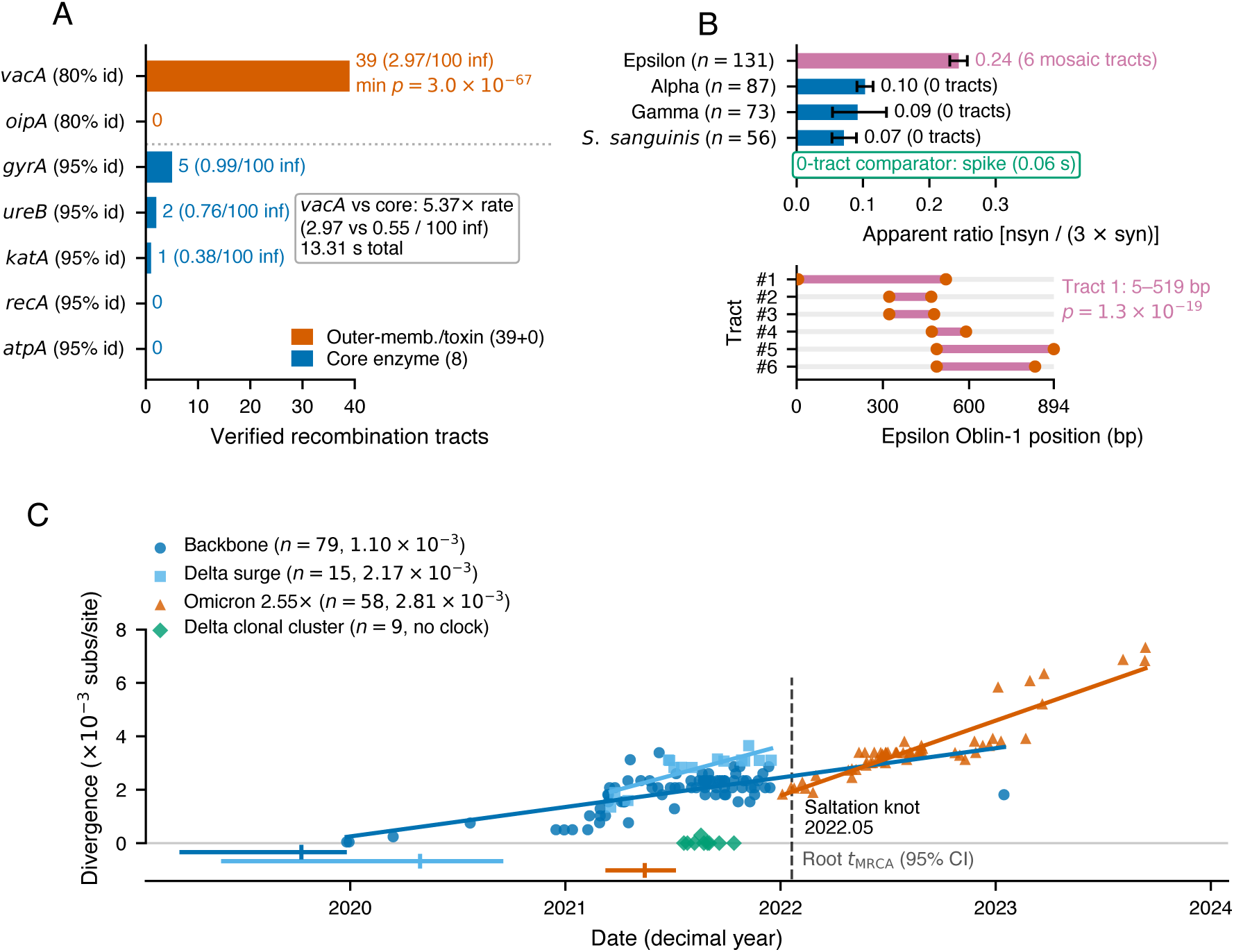
Recombination cartography (RhizAeon v0.1.0b3) and molecular clock dating (ChronAeon v0.1.0) on disassembler v0.1.0 downsampled cohorts. (A) Verified recombination tracts across seven single-copy *Helicobacter pylori* loci (*n* = 200 haplotypes per locus, *n* = 184 in strict *gyrA*, 13.3 seconds total, 80% identity gates for *vacA* and *oipA* vs. 95% for core enzymes). Cytotoxin *vacA* yields 39 tracts (2.97 per 100 informative sites) against 0 in *oipA* and 8 across two of five core enzymes (0.55 per 100 informative sites), a 5.37-fold rate enrichment (*p* = 1.0 × 10*^−^*^6^). Multi-copy adhesins (*babA*, *babB*, and *sabA*) are excluded because paralog chimeras mimic recombination. (B) Upper bars compare apparent Oblin-1 nsyn*/*(3 syn) ratios with 95% sequence-bootstrap confidence intervals across Epsilon (*n* = 131, 6 tracts), Alpha (*n* = 87), Gamma (*n* = 73), and *Streptococcus sanguinis* obelisks (*n* = 56), with the zero-tract SARS-CoV-2 spike comparator (0.06 seconds) boxed below. The lower track maps tracts #1–#6 along the 894-bp Epsilon Oblin-1 locus. (C) Unsupervised ChronAeon AutoClock partitioning across *n* = 161 dated SARS-CoV-2 spike sequences (2019.99–2023.70, four regimes), plotting divergence against date alongside the 2022.05 saltation shift and horizontal 95% root-date confidence intervals plotted below *y* = 0. Vermilion marks *vacA*, breakpoints, and Omicron, dark blue marks core enzymes, zero-tract obelisks, and the backbone, sky blue marks the Delta surge, bluish green marks comparators and the Delta clonal cluster, and reddish purple marks Epsilon tracts. Abbreviations: CI, confidence interval, bp, base pairs, and subs, substitutions.

### Obelisks

Obelisks are tiny, apparently circular RNAs, roughly 1,000 nucleotides long, recently discovered in human gut and oral metatranscriptomes, with one lineage shown to replicate in the oral bacterium *Streptococcus sanguinis* (Zheludev et al., 2024). Unlike plant viroids, which encode no proteins, obelisks carry a conserved protein called Oblin-1, hypothesized to bind RNA, that occupies most of the genome, and some lineages carry a smaller second protein called Oblin-2 alongside folded non-coding regions. Because obelisks are known almost entirely from complex human metatranscriptomes rather than from isolates, they are invisible to standard metadata queries and ideal for the kmindex and walker pathway.

We searched 141 microbiome kmindex shards (62 bacterial metagenomic, 28 human metagenomic, 29 bacterial metatranscriptomic, and 22 human metatranscriptomic shards at score threshold 0.20, *z* = 6) using five representative obelisk lineages (Alpha, Beta, Gamma, Epsilon, and an oral *Streptococcus sanguinis* lineage) and ran walker across 924 unique candidate SRA runs (Table 1). In 27.4 minutes, walker recovered 350 complete, unclipped full-length consensus sequences (311 distinct sequences) alongside 931 minor within-sample variant paths. Only three Beta primaries were recovered, so downstream comparisons focused on the four remaining clades. The original description also flags the Gamma genome as possibly chimeric (Zheludev et al., 2024), so we treat Gamma constraint estimates as provisional.

Comparing evolutionary patterns across the four large obelisk clades revealed elevated apparent amino-acid diversity and mosaic structure in the Epsilon lineage (Figure 4B). In Alpha (*n* = 87), Gamma (*n* = 73), and oral *S. sanguinis* (*n* = 56), Oblin-1 showed low apparent nonsynonymous-to-synonymous ratios (nsyn*/*(3 syn) = 0.103 [95% bootstrap CI 0.091–0.115], 0.092 [0.054–0.135], and 0.071 [0.053–0.090]). Those three clades also showed minimal epistasis (3 of 630 pairs in Alpha Oblin-1 and 0 in Alpha Oblin-2) and zero recombination tracts in RhizAeon. Epsilon Oblin-1 (*n* = 131) reached more than double that ratio at 0.244 (95% CI 0.231–0.257, covering 1,912 nonsynonymous and 2,611 synonymous differences). Supplied with external FastTree topologies, HyphAeon found zero codons passing Benjamini–Hochberg correction in any obelisk alignment (12 of 121 variable codons in Epsilon Oblin-1 and 3 of 36 in Alpha Oblin-1 at nominal *p* ≤ 0.05), yet flagged 108 of 7,260 co-evolving pairs across four linked sectors in Epsilon Oblin-1. Running RhizAeon took 0.89 seconds on Epsilon Oblin-1 and mapped six mosaic recombination tracts spanning hundreds of bases, against zero tracts in the other three clades and in a non-Oblin-1 SARS-CoV-2 spike zero-tract comparator (0.06 seconds).

Our v0.2.0 walker analysis, however, adds an essential biological caveat to that mosaic signal. Across the full-harvest run, walker flagged two-allele mixtures consistent with co-colonization, near a 2:1 abundance ratio, in 57.4% of walked runs (201 of 350 walks across 340 distinct accessions, or 347 of 594 across all ten walker runs). Mixture rates reached 61.1% (80 of 131) in Epsilon, 42.0% (37 of 88) in Alpha, 61.4% (43 of 70) in Gamma, and 69.0% (40 of 58) in *S. sanguinis*. Co-infecting strains inside the same microbiome provide the physical setting for RNA template switching (Jackson et al., 2021), yet collapsing two co-existing genotypes into a single consensus sequence can also mimic mosaic junctions, and simulation benchmarking against mixed-strain chimeras was not conducted here (Kosakovsky Pond et al., 2026a). Because Gamma and *S. sanguinis* carry comparable or higher mixture rates yet yield zero tracts, mixture frequency alone does not generate the Epsilon tracts, which remain a hypothesis awaiting phase-aware Aeon reruns.

Obelisks also show that our pipeline handles non-coding RNAs alongside overlapping shadow frames. Because walker traverses raw nucleotide assembly graphs without assuming a triplet reading frame, it reconstructs full circular obelisk RNAs across coding and non-coding structural regions alike. Likewise, tiler automatically switches off codon checks when a query is non-coding. On obelisk alignments, automatic strand orientation and coverage filtering separated intact coding Oblin-2 frames on the sense strand (Alpha and Gamma) from antisense shadow regions. In those antisense shadow frames, Epsilon carried 123 internal stops across 88 codons in 71 of 131 sequences (1.07 stops per 100 codons), and *S. sanguinis* carried 17 stops across 68 codons in 16 of 42 sequences (0.60 stops per 100 codons).

### Helicobacter pylori

To test the LexicMap and tiler pathway on a freely recombining bacterial pathogen (Suerbaum et al., 1998; Falush et al., 2001), we examined *Helicobacter pylori*, the chronic human stomach colonizer responsible for peptic ulcers and gastric cancer (Tomb et al., 1997). Across decades of persistent infection, *H. pylori* conserves core metabolic enzymes while remodeling surface adhesins and secreted toxins to evade host immunity (Atherton et al., 1995; Alm et al., 2000). We queried 20 conserved and virulence genes at a strict 95% sequence-identity gate alongside five hypervariable outer-membrane and toxin loci at a looser 80% identity gate, spanning thousands of assembled bacterial genomes per locus (Table 1).

Processing all 25 LexicMap gene scans through tiler took just 36 seconds and handled three bacterial edge cases cleanly. First, counting genuine in-frame deletions as covered sequence recovered 1,605 clean sequences of the fluoroquinolone target DNA gyrase (*gyrA*), preserving a three-base C-terminal deletion (2470_2472del, encoding N824del) present in 804 of 1,605 runs (212 of 462 unique haplotypes) (Tshibangu-Kabamba and Yamaoka, 2021). Second, automatic non-coding mode recovered 2,040 clean nucleotide sequences of flagellar cap locus *fliD*, avoiding false frameshift rejections on a gene annotated as a split pseudogene in the standard reference genome (Tomb et al., 1997). Third, reading-frame screening separated natural phase-variation frameshifts in outer-membrane adhesin *oipA* (372 strict and 800 loose runs carrying CT-repeat length shifts that switch translation on or off (Yamaoka et al., 2000)). Meanwhile, overlap-conflict checks flagged 4,306 multi-copy paralog collisions across the five loose scans of the *babA*, *babB*, *sabA*, *oipA*, and *vacA* families (Ilver et al., 1998; Mahdavi et al., 2002; Alm et al., 2000). In *babA* alone, this filter isolated 1,576 paralog mixes while retaining 408 clean full-length and 358 partial sequences (766 combined).

Downstream Aeon scans on downsampled panels of 200 *H. pylori* haplotypes per gene (184 deletion-free haplotypes in strict *gyrA*) completed in 135 seconds across the four HyphAeon runs (Table 2). Across the full coding sequences, neither housekeeping locus exceeded chance selection expectations (Figure 3A). Urease *ureB* (570 codons, 95.3% invariable) showed 1 of 27 variable codons at nominal *p* ≤ 0.05 against 1.35 expected by chance (0 at *q* ≤ 0.05, 1 of 351 pairs at *q* ≤ 0.05). Likewise, *gyrA* (828 codons) showed 13 of 188 variable codons at nominal *p* ≤ 0.05 against 9.40 expected (binomial *p* = 0.16, 0 at *q* ≤ 0.05, 14 of 17,578 pairs). Within *gyrA*, however, the single-site plasticity score Φ ranked quinolone-binding pocket residue Asn87 (Tshibangu-Kabamba and Yamaoka, 2021) as the single most constrained site across all 828 codons (Φ = 0.1957), varying across 37 of 184 deletion-free haplotypes (carrying Thr, Lys, Ile, and Tyr substitutions). By contrast, loose surface scans showed elevated co-selection density even with zero codons below *q* ≤ 0.05. Adhesin *oipA* reached 32 of 121 variable codons at nominal *p* ≤ 0.05 (4 at *q* ≤ 0.10) and 370 of 7,260 co-evolving pairs (5.10%) across two permutation-significant sectors (*C* = 0.86 and 0.82). Cytotoxin *vacA* (Gangwer et al., 2010) reached 115 of 548 nominal codons (3 at *q* ≤ 0.10) and 5,185 of 149,878 pairs (3.46%). Because tiler removes phase-variation frameshifts before alignment, the *oipA* panel samples intact in-frame alleles without out-of-frame switched states.

Adding RhizAeon separated recombination from point-mutation co-evolution in 13.3 seconds across seven single-copy loci (Figure 4A). Excluding the multi-copy adhesin families (*babA*, *babB*, and *sabA*), cytotoxin *vacA* (80% identity) carried 39 verified recombination tracts (2.97 per 100 informative sites). Five 95%-identity core enzymes combined yielded 8 tracts across two of five loci (0.55 per 100 informative sites: 5 in *gyrA*, 2 in *ureB*, 1 in *katA*, and 0 in *recA* and *atpA*). Thus, *vacA* shows a 5.37-fold enrichment per informative site (*p* = 1.0 × 10^−6^). Both loose surface cohorts retain strong continuity with v0.2.0: 81.0% of *vacA* (162 of 200) and 73.0% of *oipA* (146 of 200) haplotypes match v0.2.0 clean sequences identically, whereas those 39 tracts warn that *vacA*’s 5,185 co-evolving pairs reflect extensive domain shuffling (Atherton et al., 1995; Gangwer et al., 2010) alongside direct residue coupling. Conversely, *oipA* yielded zero tracts across 275 informative sites, pointing to point-mutation epistasis instead of long-tract exchange.

### Practical Guidelines and Limitations

Taken together, these three case studies yield clear practical rules for archive-scale evolutionary mining. When studying close relatives at low sequence divergence, uncultured RNAs in complex metagenomes, non-coding loci, or intra-sample mixtures, pair kmindex with walker because unitig graphs preserve local variants that fragment contigs. When screening millions of assembled viral or bacterial datasets across broader divergence, pair LexicMap with tiler, setting a strict 95% identity threshold for conserved housekeeping genes and looser 80% gates for hypervariable surface or toxin genes while relying on overlap-conflict checks to isolate multi-copy paralogs. Together, these three systems show that metadata-free cohort assembly uncovers shared evolutionary regimes across RNA viruses, uncultured circular RNAs, and recombining bacterial pathogens, distinguishing point-mutation epistasis from domain shuffling and saltation clock shifts from clonal clusters.

Several methodological boundaries and unverified steps remain open. First, exact 21-mer matching in walker fails abruptly below 70% query containment, and single-reference searches miss distant homologs absent from the query sequence (reference-ascertainment bias). Second, neither walker nor tiler phases full-gene haplotypes across reads: tiler emits one consensus per run, walker writes fork alleles separately, and co-colonizing strains in metagenomes can still create chimeric consensus sequences such as the mixed Epsilon obelisk primaries flagged above. Third, all Aeon evaluations in this paper ran on downsampled panels of at most 200 taxa conditioned under disassembler v0.1.0, and date granularity for uncalibrated SRA runs occasionally falls back to month-midpoint or public-release dates. Native multi-thousand-sequence Aeon benchmarks, head-to-head runtime and concordance comparisons against classical tools (MEME, BUSTED, GARD, TreeTime, and LSD2) on identical alignments, simulation checks on mixed-consensus false positives, and reruns on v0.2.0 deletion-rescued cohorts remain essential next steps. Finally, the public archive itself shifts continuously, expanding by 3.1% between our v0.1.0 and v0.2.0 SARS-CoV-2 queries.

Routine archive-wide comparative genomics also rests on accessible infrastructure and rigorous engineering. Both kmindex and LexicMap run on public Galaxy servers (The Galaxy Community, 2024), freeing biologists from terabyte storage overhead, while six Galaxy wrappers for disassembler are under community review and the CLI toolchain runs on a single compute node. Verifiable agentic development practices (Baker et al., 2026), checksum audits, and golden test fixtures caught real biological traps before release—including internal viral deletions, terminal read-clipping biases, and the *H. pylori fliD* pseudogene—making archive-scale cohort assembly reproducible for everyday evolutionary work.

## Methods

### Unitigs

When short sequencing reads are split into overlapping fixed-length words (*k*-mers) and linked whenever adjacent words overlap by *k* − 1 bases, they form a de Bruijn assembly graph (Compeau et al., 2011). A unitig is a maximal non-branching path through that graph, where every internal node has one incoming edge and one outgoing edge. Whenever true sequence variation, mixed strains, repeats, or sequencing errors create a fork or bubble, the path stops and a new unitig begins. Because unitigs never collapse across branches, they preserve the *k*-mer graph near-losslessly: every *k*-mer above a low abundance threshold is retained, so alleles, minor intra-sample variants, and structural junctions present in the raw reads stay intact alongside their mean *k*-mer read depth.

In Logan, compacted de Bruijn unitigs (*k* = 31, 30-bp overlaps, and per-unitig *k*-mer abundance tags) were built per SRA run with BCALM2 (Kodama et al., 2012; Chikhi et al., 2016, 2025). The initial Logan v1 freeze assembled 27.3 million unitig runs (96.0% of archive volume, compressing 48.2 petabases of raw reads to 2.18 petabytes). The expanded v1.2 freeze spans 38.1 million unitig runs (99.5% of volume, compressing 86.6 petabases of raw reads into 8.49 petabases uncompressed and 4.19 petabytes compressed) (Chikhi et al., 2025; Faure et al., 2026).

### Contigs

A contig is a longer linear sequence built by simplifying the unitig graph and selecting one consensus path across small branches. During graph simplification, an assembler removes short dead-end tips caused by sequencing errors, collapses small low-depth bubbles into the dominant allele, and merges the surviving paths into extended sequences. That simplification drops minor intra-sample alleles and low-abundance branches, yet yields longer, cleaner sequences that compress far more tightly than unitigs and support direct seed-and-extend alignment across divergent species.

In Logan, contigs (≥ 150 bp) were assembled per SRA run with minia3 (Chikhi et al., 2025). The initial v1 freeze assembled 26.8 million contig runs (385 terabytes compressed, revised to 315 terabytes in v1.1 after two assembler defects were corrected). The expanded v1.2 freeze spans 37,377,661 contig runs (92.3% of archive volume, 1.80 petabases uncompressed and 623 terabytes compressed) (Chikhi et al., 2025; Faure et al., 2026).

### Archive-Wide Sequence Indices: Inverted Bloom Filter Matrices and Lexico-graphic Seed Tables

To make archive-wide unitig search feasible, kmindex stores canonical *k*-mer presence in inverted Bloom filter matrices (Bloom, 1970), an approximate membership query data structure constructed with kmtricks (Lemane et al., 2022, 2024; Chikhi and Rizk, 2013). Each sequencing run is first summarized by a Bloom filter, a bit array whose positions are set by hashing every canonical *s*-mer present in that run’s Logan unitigs. Instead of scanning millions of run filters sequentially, kmtricks transposes those bit arrays into an inverted matrix where each hash row maps straight across every indexed run in a shard. At query time, kmindex applies the findere algorithm with extension parameter *z* = 6 (Robidou and Peterlongo, 2021). Instead of accepting single indexed 25-mers (*s* = 25) on their own, findere tests extended 31-mers (*k* = *s* + *z* = 31) by requiring all *z* + 1 = 7 consecutive overlapping 25-mers inside each 31-mer to test positive. That requirement cuts the Bloom filter false-positive rate exponentially to roughly 0.005% without enlarging the index. Across the hosted usegalaxy.org Logan unitig catalog (Lemane et al., 2024), this index covers approximately 2 × 10^15^ canonical *k*-mers across 23.4 to 27.3 million SRA runs and 45,412 reference genomes. Because a single global index would exceed memory limits, kmindex divides the catalog by sequencing strategy and broad taxonomic group into 109 named sub-indices, each further split into fixed-capacity query shards (2,869 shards total) so searches stream in parallel across categories such as viral or human microbiome datasets.

To support alignment across deeper sequence divergence, LexicMap builds a hierarchical lexico-graphic seed index over Logan contigs (Shen et al., 2026). Using lexicographic sketching, LexicMap guarantees no seed desert of 100 bp or longer (with a median spacing near one seed per 250 bp across genomes) and stores compact 31-bp seeds in a sorted prefix-and-suffix table. Upstream defaults require at least a 15-bp common prefix or suffix in a multi-seed chain (or 17 bp for a single anchor), whereas the public Galaxy wrapper raises these defaults to 17 bp and 19 bp. Matching seeds are chained into candidate regions and extended by banded local alignment into BLAST-like alignments. On usegalaxy.org, the pre-built LexicMap catalog spans 25 SRA contig categories totaling more than 340 terabytes.

### Running kmindex and LexicMap on Galaxy (usegalaxy.org)

Both search engines run through a web browser on https://usegalaxy.org without local index storage or command-line setup (The Galaxy Community, 2024). Biologists sign in to usegalaxy.org, open a history, and upload nucleotide FASTA queries with concise single-word headers using the Upload Data tool. Both search tools sit under the Assembly section of the left tool panel or via the tool search bar.

To query Logan unitigs, select kmindex query (cohorts reported here used version 0.7.0+galaxy0, and 0.7.1+galaxy0 is currently installed on usegalaxy.org). Under Kmindex source, choose Locally installed kmindex indexes and pick either taxon-specific partitions (such as viral RNA or human microbiome sub-indices) or all 109 SRA sub-indices covering genomic, metagenomic, transcriptomic, metatranscriptomic, single-cell, synthetic, and viral RNA categories. Assign the uploaded FASTA under Query sequences, keep the default Z-value of 6 to enforce findere filtering, set the Shared k-mers threshold according to the desired recall (for example, 0.20 for microbiome obelisks or 0.30 for broad viral cohorts), choose JSON output (optionally merging per-shard outputs), and launch the job. Download the JSON output (or fetch it via the Galaxy API) and pass it straight into disassembler harvest or disassembler from-kmindex.

To query Logan contigs, open LexicMap Search (cohorts reported here used version 0.9.0+galaxy5, and 0.10.0+galaxy0 is currently installed on usegalaxy.org). Select the uploaded FASTA under LexicMap query file, set LexicMap index source to Locally installed LexicMap indexes, and pick one or more of the 25 SRA contig categories. Set the top-N genome cap to 0 to return every matching accession, keep Output all columns enabled so tiler receives full coordinates and aligned sequences, and enable gzip compression for large tables. Notice that LexicMap Search defaults to 70.0% minimum identity (with 50-bp minimum alignment length, 17-bp chained and 19-bp single seed prefixes, 20-bp maximum gap, and E-value cutoff 10.0), so queries targeting disassembler tiler’s permissive 60.0% gate must lower the search identity floor to 60.0% before launching. Pass the downloaded table and query FASTA into disassembler lexicmap. Six Galaxy wrappers (suite_disassembler) are under review in tools-iuc, and the CLI installs from GitHub (https://github.com/nekrut/disassembler). Table 3 lists the primary queries, with full *H. pylori* coordinates in Supplementary Table S2.

**Table 3:** Primary reference queries, accessions, and coordinate frames used across all three case studies and calibration checks. Genetic Code Table 11 shares all sense and stop codons with Standard Code Table 1 while accommodating bacterial alternative initiation codons such as GTG in Obelisk Alpha Oblin-1.

| System / role | Reference accession | Length | Query scope and coordinate note |
|---|---|---|---|
| SARS-CoV-2 Spike | NC_045512.2 | 3,822 nt | Wuhan-Hu-1 S coding sequence (bases 21,563–25,384, 1,274 codons including terminal stop) |
| Microbiome obelisks | Obelisk_000001..000004, Obelisk_000006 | 1,023–1,182 nt | Five clade centroids from Zheludev et al. (2024) covering Alpha (000001, GTG start), Beta (000002), *S. sanguinis* (000003), Gamma (000004), and Epsilon (000006) |
| *Helicobacter pylori* | NC_000915.1 | 25 queries | Type strain 26695 markers (full HP_RS\* locus tags, WP_\* proteins, and genome coordinates in Table S2, including split pseudogene *fliD* spanning 2,057 nt) |
| HIV polymerase calibration | SRR8209082 | Pol locus | Clinical 14,500× human immunodeficiency virus run used to calibrate two-stage unitig fork pruning |

### Unitig Graph Path-Walking and Intra-Sample Variants (walker)

Local de Bruijn graph reconstruction ran in Rust via walker (the logan-walker binary, disassembler v0.2.0). Compressed BCALM2 unitig files—whose mean *k*-mer abundance tags saturate at 25,000.0 and emit an explicit saturation indicator—streamed directly from cloud object storage into an in-memory 2-bit compressed sparse row graph. Each fetch logged object byte counts, SHA-256 digests, and entity tags, which detected 3.1% upstream object churn across 5,000 sampled unitig keys under cache revalidation. Candidate graphs were anchored by exact canonical 21-mer containment requiring at least three shared 21-mers. Across 121,500 simulation trials (4,860 parameter cells) against a 4.64-megabase *Escherichia coli* unitig graph, canonical *k* = 21 anchoring yielded zero false-positive anchors while lifting gene recovery at 85% nucleotide identity from 24% (*k* = 31) to 100%.

Around verified anchor unitigs, a 5-hop breadth-first traversal extracted a local subgraph capped at 1,000,000 nodes and 4,096 MB compressed object size (350 MB in high-throughput SARS-CoV-2 waves), isolating edgeless subgraphs and malformed upstream objects cleanly via exit code 4 without halting batch runs. Reference queries aligned to extracted subgraphs using pinned GraphAligner v1.0.17b in single-threaded de Bruijn graph mode (Rautiainen and Marschall, 2020), which eliminates 1–3-nt 3^′^ terminal clipping present in releases 1.0.19 and 1.0.20. Walks had to cover at least 85% of the reference span (ref_cov) so authentic in-frame deletions passed coverage checks without penalty. When multiple reference anchors were provided, anchors ranked by non-ambiguous matched bases (subtracting N and degenerate IUPAC positions), breaking ties within 0.002 by mean unitig route depth.

Because unpruned graph alignment on clinical human immunodeficiency virus polymerase run SRR8209082 (14,500× depth) picked reference-matching minority branches at 49 of 50 discordant forks (98%, discarding majority alleles carrying a median 778-fold depth advantage), walker applies a two-stage fork filter. First, unitig branches below 1% of mean path depth are pruned (guarded by a 3-bp reference-span check) before realignment, cutting sub-1% noise alternatives by 89.4% (from 94 down to 10 across 187 walked panel runs, where 93.9% of retained alternatives exceed 5% depth). Second, at remaining biallelic forks with abundance ≥ 5.0, minor variant fraction ≥ 0.10, and indel length ≤ 30 bp, majority-allele swapping places the higher-depth allele onto the primary consensus haplotype, lowering forks where an alternative exceeds primary depth from 19.0% to 0.3%. Each walk records the minimum primary-allele fraction across two-allele sites (min_site_vaf = 1 − MAF), up to 32 deduplicated alternative bubble paths, a mixed=1 sample indicator, and sparse biallelic variant tables (snvs.tsv) queryable in DuckDB (Raasveldt and Mühleisen, 2019).

### Contig Coordinate Tiling, Conflict Filtering, and Genetic Codes (tiler)

Tabular LexicMap high-scoring segment pairs were projected onto reference coordinates using tiler (invoked as disassembler lexicmap in v0.2.0, with a disassembler tile alias planned). Qualifying segments meeting minimum nucleotide identity (60.0% permissive default, 80.0% loose bacterial surface mode, or 95.0% strict mode) entered six mutually exclusive quality categories, checking reference coverage before reading frame so short fragments entered Low-Coverage Fragment instead of inflating frameshift counts. Clean Full-Length required ≥ 80% reference coverage by default (100% in strict mode), Clean Partial spanned [50%, 80%) by default ([50%, 100%) strict), and failing runs entered Low-Coverage Fragment (*<* 50%), Flagged Premature Stops (≥ 1 internal termination codon), Flagged Frameshift, or Flagged Mixed Overlaps.

Multi-segment coordinate overlaps resolved at each reference nucleotide position *p* by interior depth min(*p* − start, end − 1 − *p*), breaking ties by segment nucleotide identity. Selecting interior bases prevented short 100%-identity wild-type bubble scraps from overwriting central mutations, eliminating 37 to 46 wrong bases per run seen under whole-segment identity ranking. Indels located within 10 bp of the ends of every carrying segment were flagged as untrusted edge artifacts and blocked from masking overlapping bases, while trusted internal deletions counted as covered reference positions.

Reading-frame, conflict, and translation checks ran uniformly across loci. Runs carrying an out-of-frame offset (Δ ≢ 0 (mod 3)) spanning at least 30 reference bases or reaching the 3^′^ gene boundary entered Flagged Frameshift. Runs where two segments placed discordant bases or indels at least 10 bp inside both boundaries entered Flagged Mixed Overlaps whenever either segment spanned at least min(100 bp, ⌊*L*_ref_ */*10⌋). That 100-bp span floor suppresses terminal-scrap false alarms while preserving 100% (1,576/1,576) of true paralog collisions in loose *H. pylori babA* scans (4,306 mixed runs across five loose outer-membrane loci versus 58 across 20 strict loci). References whose length is not divisible by three, or runs invoked in explicit non-coding mode, automatically disabled stop and frame checks so pseudogene-anchored or non-coding nucleotide alignments passed cleanly. Clean sequences collapsed online into unique haplotypes in memory proportional to distinct haplotype count while recording haplotype-distribution Shannon entropy in bits (*H* = − Σ_h_ *p_h_* log_2_ *p_h_*). Coding loci used NCBI Genetic Code Table 11, which matches Standard Code Table 1 across all internal sense codons and TAA, TAG, and TGA stop codons. Pre-release v0.2.0 runs executed from clean source commits whose JSON headers retained the pre-tag version string 0.1.1 alongside exact commit hashes.

### Diversity Sieving and Frame-Verified Codon Alignment

Clonal redundancy was thinned in disassembler sieve, wrapping kmer-diversity-sieve v0.2.0 (Broder, 1997; Ondov et al., 2016). Sequences were hashed into canonical 15-mer MinHash sketches across 16 locality-sensitive hashing bands (sketch size 256) and evaluated in deterministic input order. A candidate sequence *S_i_* was promoted only when its Jaccard novelty against every already-promoted representative *S_j_*satisfied:

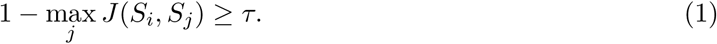

Whole-genome defaults in the standalone binary (*τ* = 0.15, sketch size 64, minimum length 27,000 bp) compress 17.6 million SARS-CoV-2 genomes into 1,443 representatives in roughly 20 minutes across 128 ranks, yet collapse single genes to one row. Wrapper single-gene defaults set *τ* = 0.005 (roughly 0.017% nucleotide divergence at *k* = 15), byte-exact pre-deduplication, minimum length fraction 0.85 for unitig walks and 0.95 for contig cohorts, and gap ceiling 0.05 (or 1 − coverage gate + 0.05). Under strict Spike tiler inputs (114,293 sequences including the ancestral reference, 0 short and 0 gappy), the sieve suppressed 195 exact duplicates and 58,375 MinHash-similar sequences to promote 55,723 representatives. Permissive Spike inputs (253,988 sequences) yielded 238 exact duplicates and 130,574 MinHash-suppressed sequences at a 25% gap cap (123,176 promoted), compared with 86,100 sequences exceeding a 5% gap cap and 88,395 MinHash-suppressed sequences at 5% (79,255 promoted). Version 0.2.0 dynamically resizes its band hash table and *k*-mer buffer—eliminating prior 65,536-record and 65.5-kb caps—and disables warmup outlier rejection so zero divergent haplotypes are dropped on admission, preserving Robinson-Foulds distance 0 on Zika virus NS5 and retaining all 17 clinical HIV polymerase walks (2%–3% divergence).

Conditioned sequences aligned in disassembler msa using MAFFT v7.526 (--auto --keeplength --preservecase --addfragments, or --aligned pass-through for pre-tiled contig cohorts) (Katoh and Standley, 2013), paired with approximate maximum-likelihood trees from FastTree (-nt -gtr) (Price et al., 2010) when supplied externally. HyphAeon does not require a tree as input: given none, it constructs an internal neighbor-joining tree from the alignment. The three obelisk HyphAeon runs (Alpha Oblin-1, Alpha Oblin-2, and Epsilon Oblin-1) received FastTree topologies, whereas all *H. pylori* and SARS-CoV-2 spike runs used the internal neighbor-joining tree. A dual 70% reference-coverage filter (applied before and after fragment addition) plus automatic 15-mer and 11-mer reverse-complement strand orientation removed overhanging gap fragments and reversed strands. Stratified kmindex sampling closed the upper match interval at [0.90, 1.00] inclusive so exact-match runs were retained.

### Empirical Case-Study Specifications

SARS-CoV-2 spike contig tiling processed 5,878,944 LexicMap segments across 4,987,809 SRA runs (3,822-bp ancestral reference NC_045512.2) under strict (≥ 95% identity) and permissive (≥ 80% coverage, ≥ 60% identity) gates. Sieved counts include the kept reference: 55,723 strict representatives (5.5 minutes), 123,176 permissive representatives under a 25% gap cap (16.5 minutes), and 79,255 under a 5% gap cap (11.3 minutes), executed on a local workstation. Strict tiling finished in 78 minutes on a 32-core virtual machine (122 GB RAM, 58 GB disk). Permissive tiling took 282 minutes on a 64-core server (nn1) sharing resources with concurrent 48-worker unitig walks and writing a 25-GB cohort ledger. On nn1, the ancestral kmindex-to-walker wave drew 20,000 stratified runs (13,333 in [0.90, 1.00], 4,000 in [0.70, 0.90), and 2,667 in [0.30, 0.70)) from 5,138,371 viral-shard hits. Streaming 125 GB across 19,892 fetched unitig files took 2 hours 38 minutes (158 minutes total) to reconstruct 11,881 primaries (10,914 full span, 7,049 unique, 5,178 sieved excluding reference in 25 seconds, and 20,390 alternatives).

Microbiome obelisk analyses queried five clade centroids (Zheludev et al., 2024) across 141 microbiome kmindex shards (62 bacterial metagenomic, 28 human metagenomic, 29 bacterial metatranscriptomic, and 22 human metatranscriptomic shards at *z* = 6, score threshold 0.20), recovering 924 unique candidate runs across five _top100 and five _all walker runs in 27.4 minutes on the 32-core virtual machine. Full-harvest (_all) recovery yielded 350 unclipped primaries (0/350 clipped versus 34/345 in v0.1.0, 311 distinct sequences, and 931 alternatives). Genotype mixtures (mixed=1) appeared in 57.4% of full-harvest runs (201/350, median minimum primary-allele fraction 0.653: Epsilon 80/131, Alpha 37/88, Gamma 43/70, *S. sanguinis* 40/58, Beta 1/3) and 58.4% across all ten runs (347/594, median 0.658). Post-alignment 70% coverage filtering dropped 16 overhang rows (28%–56% coverage) from 58 walked *S. sanguinis* antisense *oblin2* primaries and retained 42 clean rows (16/42 carrying 17 internal stops across 68 codons, or 0.60 stops/100 codons, versus 71/131 rows carrying 123 stops across 88 codons in Epsilon antisense *oblin2*, or 1.07 stops/100 codons). Apparent nonsynonymous-to-synonymous proxies were computed as nsyn*/*(3 syn) relative to each clade reference with 2,000 sequence-bootstrap replicates: Oblin-1 reached 0.244 (95% CI 0.231–0.257, 1,912/2,611) in Epsilon, 0.103 (0.091–0.115, 311/1,006) in Alpha, 0.092 (0.054–0.135, 19/69) in Gamma, and 0.071 (0.053–0.090, 211/989) in *S. sanguinis*, alongside 0.279 (0.245–0.317, 159/190) in Alpha Oblin-2 and 0.044 in Gamma Oblin-2.

Bacterial *H. pylori* scans filtered one shared 20-locus LexicMap table (207,321 rows, 2,620 to 26,260 evaluated runs per locus) at 95% strict identity plus five loose outer-membrane/toxin queries (*babA*, *babB*, *oipA*, *sabA*, and *vacA*) at 80% identity on the local workstation in 36 seconds (Table S2). Under v0.2.0, strict *gyrA* doubled from 796 to 1,605 sequences (462 haplotypes) by rescuing 2470_2472del (N824del, 804/1,605 runs, 212/462 haplotypes, 278 indel haplotypes total), and pseudogene-anchored *fliD* recovered 2,040 rows (up from 12) with zero false frameshifts on the 2,057-bp type-strain pseudogene. Reading-frame checks isolated 372 strict and 800 loose *oipA* CT-repeat frameshifts, loose *vacA* reached 2,059 clean full-length sequences (903 haplotypes, 68 mixed), and the five loose surface scans flagged 4,306 mixed paralog runs. In *babA*, v0.1.0 returned 1,528 full-length and 813 partial clean rows (2,341 combined), whereas v0.2.0’s 100-bp overlap filter isolated 1,576 paralog mixtures to leave 408 clean full-length and 358 partial rows (766 combined, a reduction of 1,575).

### Selection, Mutational Stasis, and Epistasis (HyphAeon)

All evolutionary analyses in the Aeon suite ran on cohort alignments from our frozen initial (v0.1.0) snapshot on an Apple Silicon workstation (macOS Darwin 25.3.0 arm64, MPS backend) and were audited for haplotype concordance against disassembler v0.2.0 clean cohorts (Table S3). Core *H. pylori* housekeeping alignments match v0.2.0 at 99.5% to 100.0% (including all 184 deletion-free strict *gyrA* haplotypes before v0.2.0 rescued 278 indel haplotypes), while single-copy *oipA* (146/200, 73.0%) and *vacA* (162/200, 81.0%) retain their clean backbones. The 161-haplotype SARS-CoV-2 spike panel (ancestral reference plus the first 160 ordered LexicMap haplotypes by isolate count across 1,274 codons, 110 variable) came from the v0.1.0 permissive cohort (254,915 clean haplotypes spanning 1,154,590 runs). Those 160 haplotypes account for 545,898 runs across 15 countries (led by the USA with 110 and the UK with 23), and 140 of them carry in-frame deletions (80.9% to 100% coverage). Each *H. pylori* Aeon panel took the first 200 ordered haplotypes from its clean v0.1.0 cohort (all 184 in strict *gyrA*), pre-filtering *oipA* phase-variation frameshifts before selection scans.

Codon selection, digital deep mutational scanning (Fowler and Fields, 2014), and residue epistasis ran in HyphAeon v0.2.0 (Kosakovsky Pond et al., 2026c), comprising 1.91 million axial transformer backbone parameters and 2.46 million parameters across all task heads, trained against MEME and BUSTED (Murrell et al., 2012, 2015). In the companion method preprint (Kosakovsky Pond et al., 2026c), HyphAeon achieved ROC-AUC up to 0.942 (mean 0.659), precision-recall lift up to 25.8-fold (mean 7.7-fold), and Spearman *ρ* up to 0.983 against MEME, though no direct comparator run was performed here. At each codon *s*, simulated single-amino-acid scanning evaluates all 19 substitutions away from the wild-type residue to measure within-gene intrinsic mutational plasticity:

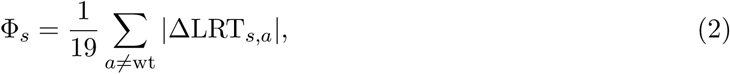

and the gene-wide summary 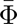 is its arithmetic mean across all codons in that alignment. Because Φ*_s_* normalizes relative to each alignment’s diversity, Φ*_s_* supports within-gene rigidity ranking instead of pooled cross-gene comparison. Co-selection networks retained pairs among the 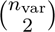 variable-codon combinations at false-discovery rate *q* ≤ 0.05 and tested spectral sector coherence *C* via 10^4^ null permutations.

Across *H. pylori*cohorts (Table S4), HyphAeon finished in 135 seconds total (18.9 to 69.1 seconds per gene). Full-gene scans of *ureB* (570 codons, 27 variable, 20.08 s, 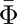 = 3.160) and *gyrA* (828 codons, 188 variable, 26.54 s, 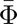 = 3.150) did not exceed chance selection expectations. Specifically, *ureB* yielded 1 of 27 variable codons at nominal *p* ≤ 0.05 against 1.35 expected (0 at *q* ≤ 0.05, 1 of 351 pairs at *q* ≤ 0.05), and *gyrA* yielded 13 of 188 nominal codons against 9.40 expected (binomial *p* = 0.16, 0 at *q* ≤ 0.05, 14 of 17,578 pairs). Within *gyrA*, single-site Φ still ranked quinolone-binding pocket residue Asn87 first in rigidity across all 828 codons (Φ = 0.1957, 37 variant haplotypes among 184). Adhesin *oipA* (306 codons, 121 variable, 18.90 s) produced 32 nominal sites (6.05 expected, 0 at *q* ≤ 0.05, 4 at *q* ≤ 0.10) and 370 of 7,260 pairs across two sectors (*p*_perm_ *<* 10^−4^: *K* = 14*, C* = 0.859 and *K* = 17*, C* = 0.818). Cytotoxin *vacA* (1,291 codons, 548 variable, 69.11 s) reached 115 nominal sites (27.4 expected, 0 at *q* ≤ 0.05, 3 at *q* ≤ 0.10) and 5,185 of 149,878 pairs. On SARS-CoV-2 spike (29.52 s), 27 of 110 variable codons cleared nominal *p* ≤ 0.05 against 5.50 expected (0 under full-CDS Benjamini–Hochberg correction, minimum *q* = 0.154). One site—Glu484 (LRT = 16.32, *p* = 1.21 × 10^−4^, 58 Omicron E484A and 5 E484K haplotypes)—survived both variable-codon Benjamini–Hochberg and Bonferroni adjustment (*q*_var_ = 0.0133). The remaining leading sites were Val445 (LRT = 8.87), Asp1118 (LRT = 8.59), Asp950 (LRT = 8.52), Ser13 (LRT = 8.42), and Pro681 (LRT = 8.35), alongside 171 of 5,995 pairs across seven sectors (4 at *p*_perm_ *<* 0.01). On obelisks, Epsilon Oblin-1 (298 codons, 121 variable, 16.08 s) yielded 12 nominal sites (0 at *q* ≤ 0.05) and 108 of 7,260 pairs across four sectors. Alpha Oblin-1 (203 codons, 12.32 s) yielded 3 of 36 nominal sites and 3 of 630 pairs, while Alpha Oblin-2 (60 codons, 5.23 s) yielded 0 of 16.

### Recombination Cartography (RhizAeon) and Molecular Clocks (ChronAeon)

We mapped intragenic mosaicism with RhizAeon v0.1.0b3 (Kosakovsky Pond et al., 2026a) across v0.1.0 alignments, identifying breakpoints from spatial shifts in cross-taxon transformer attention and testing events internally with two-sided Fisher exact tests (*p*_Fisher_) and physical clustering scores (*Z*_phys_), without running external comparators such as GARD. Across seven single-copy *H. pylori* cohorts (13.31 s total, excluding multi-copy *babA*, *babB*, and *sabA*), *vacA* (80% identity, 1,311 informative sites) carried 39 verified tracts (minimum *p* = 2.99 × 10^−67^, *Z*_phys_ = 10.52, 4.96 s, 2.97 per 100 informative sites) and *oipA* carried zero (0.57 s, 275 informative sites). By comparison, five 95%-identity core enzymes (*gyrA* 5/505, *ureB* 2/263, *katA* 1/261, *recA* 0/195, and *atpA* 0/220) totaled 8 tracts across 1,444 informative sites (0.55 per 100 informative sites). Cytotoxin *vacA* thus showed a 5.37-fold informative-site rate enrichment (*p* = 1.0 × 10^−6^, exact two-sided Poisson rate-ratio test). On obelisks, Epsilon (*n* = 131, 894 bp, 227 informative sites) returned 6 tracts in 0.89 s (Tract 1 bp 5–519, *p* = 1.28 × 10^−19^, *Z*_phys_ = 16.94). Auditing against v0.2.0 confirms 12 of 13 recombinant Epsilon isolates, 5 of 6 representatives, all 6 carrier clades, and 9 of 9 parents persist alongside 61.1% mixed=1 co-colonization. Alpha (*n* = 87, 0.22 s), Gamma (*n* = 73, 0.04 s), oral *S. sanguinis* (*n* = 56, 0.08 s), and SARS-CoV-2 spike (*n* = 161, 0.06 s) each returned zero tracts.

Unlike tree-conditioned molecular clocks (To et al., 2016; Sagulenko et al., 2018), ChronAeon v0.1.0 (Kosakovsky Pond et al., 2026b) estimated SARS-CoV-2 spike divergence directly from pairwise Tamura-Nei 1993 nucleotide distances (Tamura and Nei, 1993) across the 161 dated haplotypes (2019.99–2023.70, 2.73 s). ENA metadata dates normalized month-only entries (YYYY-MM) to the 15th midpoint and year-only entries to July 1, falling back to first_public when collection date was absent. Leave-one-out tip dating achieved an 82.8-day median absolute error (mean 97.5 days, predictive *R*^2^ = 0.714), and bilinear surge modeling placed the Omicron saltation knot at 2022.05 (ΔAIC = −50.23, *R*^2^ = 0.844). Recursive Fiedler bisection in AutoClock on the normalized sequence-affinity graph Laplacian **L**_sym_ = **I** − **D**^−1^*^/^*^2^**AD**^−1^*^/^*^2^ detected an 87.8-fold relative eigengap cliff between *K* = 4 (0.0649) and *K* = 5 (0.000739), selecting *K*^∗^ = 4 regimes in under 5 seconds (ΔAICc = −289.5). Inside each discovered community, divergence was anchored to an earliest-weighted continuous soft profile root on the nucleotide simplex with decay parameter *γ* and community baseline time *t*_0_:

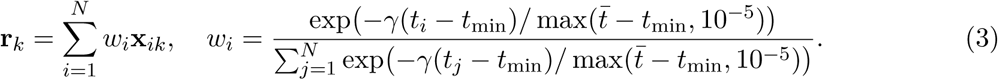

Substitution rates 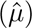, origin dates 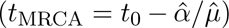, and exact analytical 95% Fieller confidence intervals 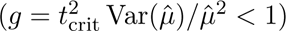 followed Fieller’s theorem (Fieller, 1954; Kosakovsky Pond et al., 2026b) (full stats in Table S4). Regime 1 captured the ancestral, Alpha, and Delta backbone (*n* = 79, 0 Omicron, *µ* = 1.10 × 10^−3^ subs/site/yr, *R*^2^ = 0.467, *t*_MRCA_ = 2019.77, CI [2019.21, 2019.98]), and Regime 2 captured a Delta surge (*n* = 15, 12 Delta + 2 Alpha + 1 D614G, *µ* = 2.17 × 10^−3^, *R*^2^ = 0.674, *t*_MRCA_ = 2020.32, CI [2019.40, 2020.71]). Regime 0 isolated Omicron (*n* = 58, 58/58 E484A+N501Y, *µ* = 2.81 × 10^−3^, *R*^2^ = 0.826, *t*_MRCA_ = 2021.37, CI [2021.19, 2021.51]), accelerating 2.55-fold over Regime 1 (delta-method 95% CI [1.87, 3.24], Wald slope difference *z* = 7.84, *p* = 4.50 × 10^−15^), while Regime 3 (*n* = 9 Delta haplotypes over 11 weeks) had no estimable positive clock (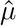 = −1.15 × 10^−4^, *p* = 0.83).

### Data Governance and Biosafety Compliance

This study consists solely of retrospective in silico computational analysis of public, de-identified nucleotide sequence records from International Nucleotide Sequence Database Collaboration repositories. No human participants were recruited, no clinical samples were collected, and no live viral, bacterial, or subviral agents were cultured or handled. No nucleic-acid synthesis, genetic modification, or gain-of-function work was performed, and all work complied with institutional biosafety policies and federal Dual Use Research of Concern requirements.

### Use of Generative AI

Generative AI tools assisted in preparing this study. Anthropic Claude models, accessed through Claude Code between September and November 2026, were used to draft and revise parts of the disassembler codebase, run and summarize cohort-conditioning and evolutionary analyses, generate figure-plotting scripts, and draft portions of the manuscript text. Commit metadata in the disassembler repository logs explicit model attribution for a subset of code contributions. Model-generated outputs entered the study only after deterministic verification against simulation benchmarks (121,500 anchor trials for *k* = 21), byte-for-byte golden fixtures, SHA-256 digests, and cross-version audits. All human authors reviewed the analyses, figures, and text and take full responsibility for every conclusion.

## Acknowledgements and Funding

Work in the authors’ groups is supported by the National Institutes of Health—specifically the National Human Genome Research Institute (U24HG002371, U24HG006620, U24HG010263), the National Cancer Institute (U24CA284167), the National Institute of Allergy and Infectious Diseases (U24AI183870), and the National Institute of General Medical Sciences (R01GM151683)—and by the National Science Foundation (grants 2419522 and 2138585).

## Author Contributions

A.C.S. and A.N. conceived the cohort-conditioning study, ran archive-scale reconstructions and provenance audits, and drafted the manuscript. R.C. advised on Logan assembly graphs and scaling, while P.S. integrated and maintained search and disassembler wrappers in public Galaxy infrastructure. D.C., S.W., and S.L.K.P. developed and ran HyphAeon, RhizAeon, ChronAeon, and kmer-diversity-sieve, and all authors revised the final manuscript.

## Data and Code Availability

Source code for disassembler (v0.2.0) is openly available at https://github.com/nekrut/disassembler, wrapping kmer-diversity-sieve (https://github.com/stevenweaver/kmer-diversity-sieve), while source code and weights for the Aeon suite reside at https://github.com/veg/HyphAeon, https://github.com/veg/RhizAeon, and https://github.com/veg/ChronAeon. Archive assembly documentation and search engines reside at Logan (https://github.com/IndexThePlanet/Logan), LexicMap (https://github.com/shenwei356/LexicMap), and https://usegalaxy.org. Conditioned cohorts, summary ledgers, and evolutionary outputs across versions 1–3 and the canonical disassembler v0.2.0 reruns are archived on Zenodo at https://doi.org/10.5281/zenodo.23737199 and https://doi.org/10.5281/zenodo.24377981 (concept record https://doi.org/10.5281/zenodo.23737198) (Seacord, 2026a,b) and mirrored at https://github.com/as042/disassembler-dump, with file-level deposit mappings in Supplementary Table S5.

## Supplementary Material

**Table S1:** Reference queries, accessions, lengths, and alignment frames used across SARS-CoV-2, microbiome obelisks, and calibration runs (see Table S2 for the 25-locus *Helicobacter pylori* query panel).

| System / clade | Reference accession | Length | Coordinate frame and notes |
|---|---|---|---|
| SARS-CoV-2 Spike | NC_045512.2 | 3,822 nt | Bases 21,563–25,384 (1,274 codons with terminal stop), query for strict and permissive LexicMap and 20,000-run kmindex stratification |
| Obelisk Alpha | Obelisk_000001_000001_000001 | 1,168 nt | Oblin-1 starts with GTG under Genetic Code Table 11 (203 codons including stop) plus tandem sense Oblin-2 (60 codons) |
| Obelisk Beta | Obelisk_000002_000001_000001 | 1,182 nt | Recovered only three primaries across 141 microbiome shards, excluded from clade-level evolutionary comparisons |
| Obelisk *S. sanguinis* | Obelisk_000003_000001_000001 | 1,137 nt | Oral *Streptococcus sanguinis* lineage (Oblin-1 231 codons with stop, antisense *oblin2* shadow region spanning 68 codons) |
| Obelisk Gamma | Obelisk_000004_000001_000001 | 1,096 nt | Reported as potentially chimeric in Zheludev et al. (2024) (Oblin-1 212 codons with stop) |
| Obelisk Epsilon | Obelisk_000006_000001_000001 | 1,023 nt | Oblin-1 298 codons with stop (894-bp coding span), antisense *oblin2* shadow region spanning 88 codons |
| HIV polymerase control | SRR8209082 | Pol locus | Clinical 14,500× depth run used to benchmark two-stage fork pruning against minor reference-matching sequencing-error branches |

**Table S2:** Complete 25-locus *Helicobacter pylori* 26695 (NC_000915.1) reference coordinates and disassembler v0.2.0 tiler cohort outcomes (20 strict scans at ≥ 95% identity and 5 loose outer-membrane/toxin scans at ≥ 80% identity, 36 seconds total). Partial clean counts exclude Clean Full-Length rows. Under v0.2.0, loose *babA* isolated 1,576 paralog mixtures from a prior 2,341-row clean pool (1,528 full + 813 partial in v0.1.0), retaining 408 full + 358 partial clean sequences.

| Locus (mode) | Locus tag | NC_000915.1 coords | Len | Runs | Full | Partial | Frame | Mixed | Haps |
|---|---|---|---|---|---|---|---|---|---|
| *atpA* (95%) | HP_RS05570 | 1196625–1198136 (c) | 1,512 | 11,552 | 2,335 | 444 | 34 | 1 | 729 |
| *babA* (95%) | HP_RS06130 | 1317838–1320039 (c) | 2,202 | 3,739 | 4 | 444 | 1 | 7 | 3 |
| *babB* (95%) | HP_RS04370 | 947580–949706 (c) | 2,127 | 3,734 | 2 | 505 | 13 | 7 | 2 |
| *cagA* (95%) | HP_RS02695 | 579921–583481 | 3,561 | 3,589 | 2 | 1,370 | 21 | 4 | 2 |
| *cagE* (95%) | HP_RS02675 | 575155–578106 (c) | 2,952 | 2,620 | 1,726 | 368 | 93 | 7 | 545 |
| *flgE* (95%) | HP_RS04245 | 920417–922573 (c) | 2,157 | 3,326 | 2,222 | 435 | 79 | 3 | 741 |
| *fliD* (95%) | HP_RS03670 | 806840–808896 | 2,057 | 3,203 | 2,040 | 376 | 0 | 2 | 660 |
| *ftsZ* (95%) | HP_RS04790 | 1042237–1043394 | 1,158 | 3,199 | 568 | 818 | 11 | 3 | 50 |
| *gyrA* (95%) | HP_RS03425 | 752512–754995 | 2,484 | 17,159 | 1,605 | 612 | 88 | 0 | 462 |
| *gyrB* (95%) | HP_RS02480 | 527686–530007 | 2,322 | 26,260 | 2,288 | 417 | 35 | 5 | 727 |
| *hpaA* (95%) | HP_RS03880 | 853957–854739 (c) | 783 | 3,167 | 2,199 | 334 | 52 | 1 | 662 |
| *katA* (95%) | HP_RS04270 | 925571–927088 (c) | 1,518 | 10,542 | 2,219 | 721 | 67 | 1 | 757 |
| *oipA* (95%) | HP_RS03145 | 684774–685691 | 918 | 3,084 | 864 | 603 | 372 | 4 | 142 |
| *recA* (95%) | HP_RS00755 | 162928–163971 | 1,044 | 7,249 | 1,973 | 819 | 29 | 1 | 466 |
| *rpsL* (95%) | HP_RS05880 | 1267881–1268288 (c) | 408 | 13,095 | 2,452 | 397 | 36 | 1 | 637 |
| *sabA* (95%) | HP_RS03530 | 779008–780924 (c) | 1,917 | 3,346 | 3 | 474 | 1 | 0 | 2 |
| *sodB* (95%) | HP_RS01915 | 398432–399073 (c) | 642 | 3,724 | 2,322 | 582 | 31 | 1 | 740 |
| *ureA* (95%) | HP_RS00370 | 77240–77956 (c) | 717 | 3,880 | 2,466 | 607 | 56 | 2 | 839 |
| *ureB* (95%) | HP_RS00365 | 75527–77236 (c) | 1,710 | 7,976 | 2,300 | 738 | 75 | 3 | 778 |
| *vacA* (95%) | HP_RS04325 | 938415–942287 | 3,873 | 3,603 | 336 | 225 | 12 | 5 | 18 |
| *babA* (80%) | HP_RS06130 | 1317838–1320039 (c) | 2,202 | 3,739 | 408 | 358 | 81 | 1,576 | 352 |
| *babB* (80%) | HP_RS04370 | 947580–949706 (c) | 2,127 | 3,734 | 386 | 427 | 293 | 1,382 | 320 |
| *oipA* (80%) | HP_RS03145 | 684774–685691 | 918 | 3,084 | 1,785 | 140 | 800 | 26 | 556 |
| *sabA* (80%) | HP_RS03530 | 779008–780924 (c) | 1,917 | 3,346 | 1,016 | 279 | 119 | 1,254 | 592 |
| *vacA* (80%) | HP_RS04325 | 938415–942287 | 3,873 | 3,603 | 2,059 | 301 | 327 | 68 | 903 |

**Table S3:** Concordance audit between the frozen disassembler v0.1.0 cohorts used for Aeon analyses and the canonical v0.2.0 cohort builds. Live unitig object churn measured across 5,000 revalidated keys was 3.1%.

| Cohort or locus | v0.1.0 snapshot | v0.2.0 canonical build | Exact concordance outcome |
|---|---|---|---|
| SARS-CoV-2 Spike (strict) | 76,360 clean sequences | 910,848 clean sequences | In-frame deletion rescue recovers 836,270 deletion-bearing runs |
| *H. pylori* core enzymes | 200 taxa per locus (184 *gyrA*) | Up to 2,335 full sequences | 99.5% to 100.0% haplotype match across *ureB*, *gyrA* (184 deletion-free), *recA*, *atpA*, *katA* |
| *H. pylori gyrA* | 796 clean (0 indels) | 1,605 clean (462 haplotypes) | All 184 deletion-free haplotypes match, 278 additional indel haplotypes rescued (2470_2472del in 804 runs) |
| *H. pylori fliD* | 12 clean sequences | 2,040 clean (660 haplotypes) | Non-coding pseudogene guard eliminates 2,481 false frameshifts |
| *H. pylori oipA* / *vacA* (loose) | 200 / 200 haplotypes | 1,785 / 2,059 clean full | 146/200 (73.0%) *oipA* and 162/200 (81.0%) *vacA* haplotypes match identically |
| Microbiome obelisks (Oblin-1) | 34/345 terminal 3′ clipped | 0/350 clipped across 5 lineages | 12/13 recombinant Epsilon isolates, 5/6 representatives, 6/6 carrier clades, and 9/9 parents persist alongside 57.4% mixed=1 |

**Table S4:** Complete statistical ledger across all HyphAeon v0.2.0, RhizAeon v0.1.0b3, and ChronAeon v0.1.0 evolutionary runs on disassembler v0.1.0 panels (≤ 200 taxa). Expected nominal counts equal 0.05 × *n*_var_. BH *q* ≤ 0.05 counts reflect full-gene adjustment (parenthetical *q*_var_ in Spike restricts BH to 110 variable codons).

| Run / locus | Taxa / cod | Var / exp | Nom *p* ≤ 0.05 | BH *q* ≤ 0.05 | Pairs (*q* ≤ 0.05) | Additional metrics (Φ̄, RhizAeon, or ChronAeon) |
|---|---|---|---|---|---|---|
| *H. pylori ureB* | 200 / 570 | 27 / 1.35 | 1 (3.7%) | 0 (0) | 1 / 351 (0.28%) | Φ̄ = 3.160, RhizAeon: 2 tracts / 263 inf (0.76/100) |
| *H. pylori gyrA* | 184 / 828 | 188 / 9.40 | 13 (6.9%) | 0 (0) | 14 / 17,578 (0.08%) | Φ̄ = 3.150 (Asn87 Φ = 0.1957), Rhiz: 5 / 505 inf (0.99/100) |
| *H. pylori oipA* | 200 / 306 | 121 / 6.05 | 32 (26.4%) | 0 (4 at 0.10) | 370 / 7,260 (5.10%) | Φ̄ = 2.547, 2 sectors (*C* = 0.859, 0.818), Rhiz: 0 / 275 inf |
| *H. pylori vacA* | 200 / 1,291 | 548 / 27.4 | 115 (21.0%) | 0 (3 at 0.10) | 5,185 / 149,878 (3.46%) | Φ̄ = 2.074, 2 sectors, Rhiz: 39 / 1,311 inf (2.97/100) |
| *H. pylori katA/recA/atpA* | 200 each | — | — | — | — | RhizAeon: 1 / 261 (*katA*), 0 / 195 (*recA*), 0 / 220 (*atpA*) |
| SARS-CoV-2 Spike | 161 / 1,274 | 110 / 5.50 | 27 (24.5%) | 0 (1 *q*<sub>var</sub>) | 171 / 5,995 (2.85%) | 7 sectors (4 at *p* < 0.01), RhizAeon: 0 / 93 inf (0.06 s) |
| Obelisk Epsilon O1 | 123 / 298 | 121 / 6.05 | 12 (9.9%) | 0 (0) | 108 / 7,260 (1.49%) | 4 sectors, RhizAeon (*n* = 131): 6 / 227 inf (Tract 1 bp 5–519) |
| Obelisk Alpha O1 / O2 | 67 / 203 (60) | 36 (16) | 3 (0) | 0 (0) | 3 / 630 (—) | RhizAeon Oblin-1: Alpha 0 / 84 inf, Gamma 0 / 12, *S. sang.* 0 / 132 |
| ChronAeon AutoClock (*K*<sup>\*</sup> = 4 regimes on 161 SARS-CoV-2 Spike haplotypes, ΔAICc = −289.5, date MAE = 82.8 days) |  |  |  |  |  |  |
| Regime 1 (Backbone) | *n* = 79 | 2019.99–2023.04 (0 Omicron: 2 D614, 20 D614G, 5 Alpha, 52 Delta), *μ̂* = 1.10 ± 0.13 × 10<sup>−3</sup>, *R*<sup>2</sup> = 0.467, *p* = 4.0 × 10<sup>−12</sup>, *t*<sub>MRCA</sub> = 2019.77 [2019.21–2019.98] |  |  |  |  |
| Regime 2 (Delta surge) | *n* = 15 | 2021.21–2021.96 (12 Delta, 2 Alpha, 1 D614G), *μ̂* = 2.17 ± 0.42 × 10<sup>−3</sup>, *R*<sup>2</sup> = 0.674, *p* = 1.7 × 10<sup>−4</sup>, *t*<sub>MRCA</sub> = 2020.32 [2019.40–2020.71] |  |  |  |  |
| Regime 0 (Omicron) | *n* = 58 | 2022.01–2023.70 (58/58 E484A+N501Y), *μ̂* = 2.81 ± 0.17 × 10<sup>−3</sup> (2.55×, 95% CI 1.87–3.24), *R*<sup>2</sup> = 0.826, *t*<sub>MRCA</sub> = 2021.37 [2021.19–2021.51] |  |  |  |  |
| Regime 3 (Delta clonal) | *n* = 9 | 2021.55–2021.78 (9/9 Delta across 11 weeks), OLS *μ̂* = −1.15 × 10<sup>−4</sup> (*R*<sup>2</sup> = 0.007, *p* = 0.828), no estimable positive clock |  |  |  |  |

**Table S5:** Provenance and reproducibility map linking manuscript figures, tables, and empirical datasets to disassembler-dump archives on Zenodo (10.5281/zenodo.23737199 and 10.5281/zenodo.24377981) and compute nodes.

| Manuscript asset | Execution host | Script / tool | Repository path in disassembler-dump |
|---|---|---|---|
| SARS-CoV-2 Spike tiler (strict / perm) | 32-core VM (122 GB RAM) / 64-core nn1 | disassembler lexicmap v0.2.0 | sars-cov-2/lexicmap/disassembler-0.2.0/ |
| SARS-CoV-2 Spike 20k walker wave | 64-core nn1 (125 GB streamed in 2h38m) | logan-walker v0.2.0 | sars-cov-2/metadata/results/walker20k_wuhan/disassembler-0.2.0/ |
| Microbiome obelisks (141 shards) | 32-core VM (58 GB disk) | logan-walker v0.2.0 (0e86dde) | obelisk/disassembler-0.2.0/ and obelisk/msa_codon/ |
| *H. pylori* 25-locus tiler panel | Local workstation (OptiPlex) | disassembler lexicmap v0.2.0 (4f8196a) | h-pylori/disassembler-0.2.0/ and h-pylori/queries/markers/ |
| Aeon evolutionary analyses (Figs 3–4, Table 2) | Apple Silicon workstation (macOS arm64) | HyphAeon v0.2.0, RhizAeon v0.1.0b3, ChronAeon v0.1.0 | showcase/PROVENANCE.md, showcase/scripts/, and showcase/results/ |
| Figures 1–4 vector and raster builds | Any workstation (Python 3.10+) | scripts/build_figures.py | Generates all four PDF/PNG/SVG figures directly from --data-dir |

## References

Richard A. Alm, James Bina, Beth M. Andrews, Peter Doig, Robert E. W. Hancock, and Trevor J. Trust. Comparative genomics of *Helicobacter pylori*: Analysis of the outer membrane protein families. Infection and Immunity, 68(7):4155–4168, 2000. doi: 10.1128/iai.68.7.4155-4168.2000.

John C. Atherton, Ping Cao, Richard M. Peek, Murali K.R. Tummuru, Martin J. Blaser, and Timothy L. Cover. Mosaicism in vacuolating cytotoxin alleles of *Helicobacter pylori*. Journal of Biological Chemistry, 270(30):17771–17777, 1995. doi: 10.1074/jbc.270.30.17771.

Dannon Baker, Clemens Blank, Dave Bouvier, Martin Čech, John Chilton, Nate Coraor, Fabio Cumbo, Jeremy Goecks, Björn Grüning, Jen Hillman-Jackson, Vahid Jalili, Delphine Larivière, Sergei L. Kosakovsky Pond, Michael C. Schatz, and Anton Nekrutenko. Verifiable agentic science on public infrastructure. bioRxiv, page 2026.10.26.761864, 2026. doi: 10.64898/2026.10.26.761864.

Burton H. Bloom. Space/time trade-offs in hash coding with allowable errors. Communications of the ACM, 13(7):422–426, 1970. doi: 10.1145/362686.362692.

Remco Bouckaert, Timothy G. Vaughan, Joëlle Barido-Sottani, Sebastián Duchêne, Mathieu Fourment, Alexandra Gavryushkina, Joseph Heled, Graham Jones, Denise Kühnert, Nicola De Maio, Michael Matschiner, Fábio K. Mendes, Nicola F. Müller, Huw A. Ogilvie, Louis du Plessis, Alex Popinga, Andrew Rambaut, David Rasmussen, Igor Siveroni, Marc A. Suchard, Chieh-Hsi Wu, Dong Xie, Chi Zhang, Tanja Stadler, and Alexei J. Drummond. BEAST 2.5: An advanced software platform for Bayesian evolutionary analysis. PLoS Computational Biology, 15 (4):e1006650, 2019. doi: 10.1371/journal.pcbi.1006650.

Andrei Z. Broder. On the resemblance and containment of documents. In Proceedings of Compression and Complexity of Sequences 1997, pages 21–29, 1997. doi: 10.1109/SEQUEN.1997.666900.

Yunlong Cao, Fanchong Jian, Jing Wang, Yuanling Yu, Weiliang Song, Ayijiang Yisimayi, Jing Wang, Ran An, Xiaosu Chen, Na Zhang, Yao Wang, Peng Wang, et al. Imprinted SARS-CoV-2 humoral immunity induces convergent Omicron RBD evolution. Nature, 614(7948):521–529, 2023. doi: 10.1038/s41586-022-05644-7.

Rayan Chikhi and Guillaume Rizk. Space-efficient and exact de Bruijn graph representation based on a Bloom filter. Algorithms for Molecular Biology, 8(1):22, 2013. doi: 10.1186/1748-7188-8-22.

Rayan Chikhi, Antoine Limasset, and Paul Medvedev. Compacting de Bruijn graphs from sequencing data quickly and in low memory. Bioinformatics, 32(12):i201–i208, 2016. doi: 10.1093/bioinformatics/btw279.

Rayan Chikhi, Téo Lemane, Raphaël Loll-Krippleber, Merce Montoliu-Nerin, Brice Raffestin, Antonio Pedro Camargo, Carson J. Miller, Mateus Bernabe Fiamenghi, Daniel Paiva Agustinho, Sina Majidian, Greg Autric, Maxime Hugues, Junkyoung Lee, Roland Faure, Kristen D. Curry, Gaetan Benoit, Yoshihiro Shibuya, Jorge A. Moura de Sousa, Eduardo P. C. Rocha, David Koslicki, Paul Medvedev, Daniel Gautheret, Purav Gupta, Thomas Quigley, Dennis Zhu, Katharina Rothe, Jessica Shen, Alejandro Morales-Tapia, Benjamin J. Buchfink, Kate Sihuta, Peter J. Roy, Grant W. Brown, Anton Nekrutenko, Robert C. Edgar, Anton Korobeynikov, Martin Steinegger, Caleb A. Lareau, Pierre Peterlongo, and Artem Babaian. Logan: Planetary-scale genome assembly surveys life’s diversity. bioRxiv, page 2024.07.30.605881, 2025. doi: 10.1101/2024.07.30.605881.

Phillip E. C. Compeau, Pavel A. Pevzner, and Glenn Tesler. How to apply de Bruijn graphs to genome assembly. Nature Biotechnology, 29(11):987–991, 2011. doi: 10.1038/nbt.2023.

Robert C. Edgar, Jeff Taylor, Victor Lin, Tomer Altman, Pierre Barbera, Dmitry Meleshko, Dan Lohr, Gherman Novakovsky, Benjamin Buchfink, Basem Al-Shayeb, Jillian F. Banfield, Marcos de la Peña, Anton Korobeynikov, Rayan Chikhi, and Artem Babaian. Petabase-scale sequence alignment catalyses viral discovery. Nature, 602(7895):142–147, 2022. doi: 10.1038/s41586-021-04332-2.

Daniel Falush, Christian Kraft, Nancy S. Taylor, Pelayo Correa, James G. Fox, Mark Achtman, and Sebastian Suerbaum. Recombination and mutation during long-term gastric colonization by *Helicobacter pylori*: Estimates of clock rates, recombination size, and minimal age. Proceedings of the National Academy of Sciences, 98(26):15056–15061, 2001. doi: 10.1073/pnas.251396098.

Roland Faure, A. H. Nakshi, Md Hasin Abrar, Rayan Chikhi, Téo Lemane, Mohammad Taheri, Seung-Hyun Won, Haonan Wu, David Koslicki, and Paul Medvedev. Sketching the SRA: Experiment-wide similarity search across 4.8 million metagenomes. bioRxiv, page 2026.10.07.757297, 2026. doi: 10.64898/2026.10.07.757297.

E. C. Fieller. Some problems in interval estimation. Journal of the Royal Statistical Society: Series B (Methodological), 16(2):175–185, 1954. doi: 10.1111/j.2517-6161.1954.tb00159.x.

Douglas M. Fowler and Stanley Fields. Deep mutational scanning: a new style of protein science. Nature Methods, 11(8):801–807, 2014. doi: 10.1038/nmeth.3027.

Kelly A. Gangwer, Carrie L. Shaffer, Sebastian Suerbaum, D. Borden Lacy, Timothy L. Cover, and Seth R. Bordenstein. Molecular evolution of the *Helicobacter pylori* vacuolating toxin gene *vacA*. Journal of Bacteriology, 192(23):6126–6135, 2010. doi: 10.1128/jb.01081-10.

Rafael S. Gonçalves and Mark A. Musen. The variable quality of metadata about biological samples used in biomedical experiments. Scientific Data, 6(1):190021, 2019. doi: 10.1038/sdata.2019.21.

Allison J. Greaney, Tyler N. Starr, Christopher O. Barnes, Yiska Weisblum, Fabian Schmidt, Marina Caskey, Michel C. Nussenzweig, Pamela J. Bjorkman, Paul D. Bieniasz, Theodora Hatziioannou, and Jesse D. Bloom. Mapping mutations to the SARS-CoV-2 RBD that escape binding by different classes of antibodies. Nature Communications, 12(1):4196, 2021. doi: 10.1038/s41467-021-24435-8.

William T. Harvey, Alessandro M. Carabelli, Ben Jackson, Ravindra K. Gupta, Emma C. Thomson, Ewan M. Harrison, Catherine Ludden, Richard Reeve, Andrew Rambaut, COVID-19 Genomics UK (COG-UK) Consortium, Sharon J. Peacock, and David L. Robertson. SARS-CoV-2 variants, spike mutations and immune escape. Nature Reviews Microbiology, 19(7):409–424, 2021. doi: 10.1038/s41579-021-00573-0.

Yu-Ning Huang, Pooja Vinod Jaiswal, Anushka Rajes, Anushka Yadav, Dottie Yu, Fangyun Liu, Grace Scheg, Emma Shih, Grigore Boldirev, Irina Nakashidze, Aditya Sarkar, Jay Himanshu Mehta, Ke Wang, Khooshbu Kantibhai Patel, Mustafa Ali Baig Mirza, Kunali Chetan Hapani, Qiushi Peng, Ram Ayyala, Ruiwei Guo, Shaunak Kapur, Tejasvene Ramesh, Dumitru Ciorbă, Viorel Munteanu, Viorel Bostan, Mihai Dimian, Malak S. Abedalthagafi, and Serghei Mangul. The systematic assessment of completeness of public metadata accompanying omics studies in the Gene Expression Omnibus data repository. Genome Biology, 26(1):274, 2025. doi: 10.1186/s13059-025-03725-0.

Dag Ilver, Anna Arnqvist, Johan Ögren, Inga-Maria Frick, Dangeruta Kersulyte, Engin T. Incecik, Douglas E. Berg, Antonello Covacci, Lars Engstrand, and Thomas Borén. *Helicobacter pylori* adhesin binding fucosylated histo-blood group antigens revealed by retagging. Science, 279(5349): 373–377, 1998. doi: 10.1126/science.279.5349.373.

Luiz Carlos Irber, N. Tessa Pierce-Ward, and C. Titus Brown. Sourmash branchwater enables lightweight petabyte-scale sequence search. bioRxiv, 2022. doi: 10.1101/2022.11.02.514947.

Ben Jackson, Maciej F. Boni, Matthew J. Bull, Amy Colleran, Rachel M. Colquhoun, Alistair C. Darby, Sam Haldenby, Verity Hill, Anastasia Lucaci, John T. McCrone, Samuel M. Nicholls, ’Aine O’Toole, et al. Generation and transmission of interlineage recombinants in the SARS-CoV-2 pandemic. Cell, 184(20):5179–5188.e8, 2021. doi: 10.1016/j.cell.2021.08.014.

Mikhail Karasikov, Harun Mustafa, Daniel Danciu, Oleksandr Kulkov, Marc Zimmermann, Christopher Barber, Gunnar Ratsch, and Andr’e Kahles. Efficient and accurate search in petabase-scale sequence repositories. Nature, 2025. doi: 10.1038/s41586-025-09603-w.

Kazutaka Katoh and Daron M. Standley. MAFFT multiple sequence alignment software version 7: improvements in performance and usability. Molecular Biology and Evolution, 30(4):772–780, 2013. doi: 10.1093/molbev/mst010.

Yuichi Kodama, Martin Shumway, and Rasko Leinonen. The Sequence Read Archive: explosive growth of sequencing data. Nucleic Acids Research, 40(D1):D54–D56, 2012. doi: 10.1093/nar/gkr854.

Sergei L. Kosakovsky Pond, Simon D. W. Frost, and Spencer V. Muse. HyPhy: hypothesis testing using phylogenies. Bioinformatics, 21(5):676–679, 2005. doi: 10.1093/bioinformatics/bti079.

Sergei L. Kosakovsky Pond, David Posada, Michael B. Gravenor, Christopher H. Woelk, and Simon D. W. Frost. Automated phylogenetic detection of recombination using a genetic algorithm. Molecular Biology and Evolution, 23(10):1891–1901, 2006. doi: 10.1093/molbev/msl051.

Sergei L. Kosakovsky Pond, Danielle Callan, Steven Weaver, Hannah Verdonk, Jordan Zehr, Anton Nekrutenko, Taylor K. Paisie, and Darren P. Martin. Clean breaks: Attention force field deconvolution of recombination across sequence manifolds. bioRxiv, page 2026.10.19.760237, 2026a. doi: 10.64898/2026.10.19.760237.

Sergei L. Kosakovsky Pond, Darren P. Martin, Brittany Rife Magalis, Sudhir Kumar, Jordan D. Zehr, Danielle Callan, Joel O. Wertheim, Steven Weaver, and Anton Nekrutenko. No chains attached: Tree-free molecular clock dating via distance geometry and evolutionary foundation models, 2026b.

Sergei L. Kosakovsky Pond, Steven Weaver, Danielle Callan, Jordan D. Zehr, Alexander G. Lucaci, Hannah Verdonk, Avery Selberg, Gallean Brown, Maria Chikina, Nathan L. Clark, Kateryna D. Makova, Darren P. Martin, and Anton Nekrutenko. HyphAeon: Attention on evolution across deep time transforms comparative genomics. bioRxiv, page 2026.09.06.749597, 2026c. doi: 10.64898/2026.09.06.749597. URL https://www.biorxiv.org/content/10.64898/2026.09.06.749597v1.

Jun Lan, Jiwan Ge, Jinfang Yu, Sisi Shan, Huan Zhou, Shilong Fan, Qi Zhang, Xuanling Shi, Qisheng Wang, Linqi Zhang, and Xinquan Wang. Structure of the SARS-CoV-2 spike receptor-binding domain bound to the ACE2 receptor. Nature, 581(7807):215–220, 2020. doi: 10.1038/s41586-020-2180-5.

Téo Lemane, Paul Medvedev, Rayan Chikhi, and Pierre Peterlongo. kmtricks: efficient and flexible construction of Bloom filters for large sequencing data collections. Bioinformatics Advances, 2(1): vbac029, 2022. doi: 10.1093/bioadv/vbac029.

Téo Lemane, Nolan Lezzoche, Julien Lecubin, Eric Pelletier, Magali Lescot, Rayan Chikhi, and Pierre Peterlongo. Indexing and real-time user-friendly queries in terabyte-sized complex genomic datasets with kmindex and ORA. Nature Computational Science, 4(2):104–109, 2024. doi: 10.1038/s43588-024-00596-6.

Jafar Mahdavi, Berit Sondén, Marina Hurtig, Farzad O. Olfat, Lina Forsberg, Niamh Roche, Jonas Ångström, Thomas Larsson, Susann Teneberg, Karl-Anders Karlsson, Siiri Altraja, Torkel Wadström, et al. *Helicobacter pylori* SabA adhesin in persistent infection and chronic inflammation. Science, 297(5581):573–578, 2002. doi: 10.1126/science.1069076.

Benoit Morel, Pierre Barbera, Lucas Czech, Ben Bettisworth, Lukas Hübner, Sarah Lutteropp, Dora Serdari, Evangelia-Georgia Kostaki, Ioannis Mamais, Alexey M. Kozlov, Pavlos Pavlidis, Dimitrios Paraskevis, and Alexandros Stamatakis. Phylogenetic analysis of SARS-CoV-2 data is difficult. Molecular Biology and Evolution, 38(5):1777–1791, 2021. doi: 10.1093/molbev/msaa314.

Ben Murrell, Joel O. Wertheim, Sasha Moola, Tashlin Weighill, Konrad Scheffler, and Sergei L. Kosakovsky Pond. Detecting individual sites subject to episodic diversifying selection. PLoS Genetics, 8(7):e1002764, 2012. doi: 10.1371/journal.pgen.1002764.

Ben Murrell, Steven Weaver, Martin D. Smith, Joel O. Wertheim, Sasha Murrell, Anthony Aylward, Kemal Eren, Tristan Pollner, Darren P. Martin, Davey M. Smith, Konrad Scheffler, and Sergei L. Kosakovsky Pond. Gene-wide identification of episodic selection. Molecular Biology and Evolution, 32(5):1365–1371, 2015. doi: 10.1093/molbev/msv035.

Sergey Nurk, Dmitry Meleshko, Anton Korobeynikov, and Pavel A. Pevzner. metaSPAdes: a new versatile metagenomic assembler. Genome Research, 27(5):824–834, 2017. doi: 10.1101/gr.213959.116.

Brian D. Ondov, Todd J. Treangen, P’all Melsted, Adam B. Mallonee, Nicholas H. Bergman, Sergey Koren, and Adam M. Phillippy. Mash: fast genome and metagenome distance estimation using MinHash. Genome Biology, 17(1):132, 2016. doi: 10.1186/s13059-016-0997-x.

Jonathan E. Pekar, Andrew Magee, Edyth Parker, Niema Moshiri, Katherine Izhikevich, Jennifer L. Havens, Karthik Gangavarapu, Lorena M. Malpica Serrano, Alexander Crits-Christoph, Nathaniel L. Matteson, Mark Zeller, Joshua I. Levy, et al. The molecular epidemiology of multiple zoonotic origins of SARS-CoV-2. Science, 377(6609):960–966, 2022. doi: 10.1126/science.abp8337.

Morgan N. Price, Paramvir S. Dehal, and Adam P. Arkin. FastTree 2 – approximately maximum-likelihood trees for large alignments. PLoS ONE, 5(3):e9490, 2010. doi: 10.1371/journal.pone.0009490.

Mark Raasveldt and Hannes Mühleisen. DuckDB: an embeddable analytical database. In Proceedings of the 2019 International Conference on Management of Data, pages 1981–1984, 2019. doi: 10.1145/3299869.3320212.

Mikko Rautiainen and Tobias Marschall. GraphAligner: rapid and versatile sequence-to-graph alignment. Genome Biology, 21(1):253, 2020. doi: 10.1186/s13059-020-02157-2.

Lucas Robidou and Pierre Peterlongo. findere: fast and precise approximate membership query. In String Processing and Information Retrieval, pages 151–163. Springer, 2021. doi: 10.1007/978-3-030-86692-9_12.

Pavel Sagulenko, Vadim Puller, and Richard A. Neher. TreeTime: Maximum-likelihood phylodynamic analysis. Virus Evolution, 4(1):vex042, 2018. doi: 10.1093/ve/vex042.

Andrew Seacord. disassembler-dump case study data, 2026a. Version 1 archive.

Andrew Seacord. disassembler-dump case study data, 2026b. Version 4 archive, disassembler 0.2.0 reruns.

Wei Shen, John A. Lees, and Zamin Iqbal. Efficient sequence alignment against millions of prokaryotic genomes with LexicMap. Nature Biotechnology, 44(8):1383–1390, 2026. doi: 10.1038/s41587-025-02812-8.

Sergey A. Shiryev and Richa Agarwala. Indexing and searching petabase-scale nucleotide resources. Nature Methods, 21(6):994–1002, 2024. doi: 10.1038/s41592-024-02280-z.

Tyler N. Starr, Allison J. Greaney, Sarah K. Hilton, Daniel Ellis, Katharine H. D. Crawford, Adam S. Dingens, Mary Jane Navarro, John E. Bowen, M. Alejandra Tortorici, Alexandra C. Walls, Neil P. King, David Veesler, and Jesse D. Bloom. Deep mutational scanning of SARS-CoV-2 receptor binding domain reveals constraints on folding and ACE2 binding. Cell, 182(5):1295–1310.e20, 2020. doi: 10.1016/j.cell.2020.08.012.

Marc A. Suchard, Philippe Lemey, Guy Baele, Daniel L. Ayres, Alexei J. Drummond, and Andrew Rambaut. Bayesian phylogenetic and phylodynamic data integration using BEAST 1.10. Virus Evolution, 4(1):vey016, 2018. doi: 10.1093/ve/vey016.

Sebastian Suerbaum, John Maynard Smith, Khairun Bapumia, Giovanna Morelli, Noel H. Smith, Erdmute Kunstmann, Isabelle Dyrek, and Mark Achtman. Free recombination within *Helicobacter pylori*. Proceedings of the National Academy of Sciences, 95(21):12619–12624, 1998. doi: 10.1073/pnas.95.21.12619.

Koichiro Tamura and Masatoshi Nei. Estimation of the number of nucleotide substitutions in the control region of mitochondrial DNA in humans and chimpanzees. Molecular Biology and Evolution, 10(3):512–526, 1993. doi: 10.1093/oxfordjournals.molbev.a040023.

The Galaxy Community. The Galaxy platform for accessible, reproducible, and collaborative data analyses: 2024 update. Nucleic Acids Research, 52(W1):W83–W94, 2024. doi: 10.1093/nar/gkae410.

Thu-Hien To, Matthieu Jung, Samantha Lycett, and Olivier Gascuel. Fast dating using least-squares criteria and algorithms. Systematic Biology, 65(1):82–97, 2016. doi: 10.1093/sysbio/syv068.

Jean-F. Tomb, Owen White, Anthony R. Kerlavage, Rebecca A. Clayton, Granger G. Sutton, Robert D. Fleischmann, Karen A. Ketchum, Hans Peter Klenk, Steven Gill, Brian A. Dougherty, Karen Nelson, John Quackenbush, et al. The complete genome sequence of the gastric pathogen *Helicobacter pylori*. Nature, 388(6642):539–547, 1997. doi: 10.1038/41483.

Evariste Tshibangu-Kabamba and Yoshio Yamaoka. *Helicobacter pylori* infection and antibiotic resistance — from biology to clinical implications. Nature Reviews Gastroenterology & Hepatology, 18(9):613–629, 2021. doi: 10.1038/s41575-021-00449-x.

Yatish Turakhia, Bryan Thornlow, Angie S. Hinrichs, Nicola De Maio, Landen Gozashti, Robert Lanfear, David Haussler, and Russell Corbett-Detig. Ultrafast sample placement on existing trees (UShER) enables real-time phylogenetics for the SARS-CoV-2 pandemic. Nature Genetics, 53(6): 809–816, 2021. doi: 10.1038/s41588-021-00862-7.

Raquel Viana, Sikhulile Moyo, Daniel G. Amoako, Houriiyah Tegally, Cathrine Scheepers, Christian L. Althaus, Ugochukwu J. Anyaneji, Phillip A. Bester, Maciej F. Boni, Mohammed Chand, Wonderful T. Choga, Rachel Colquhoun, et al. Rapid epidemic expansion of the SARS-CoV-2 Omicron variant in southern Africa. Nature, 603(7902):679–686, 2022. doi: 10.1038/s41586-022-04411-y.

Yoshio Yamaoka, Dong H. Kwon, and David Y. Graham. A *M*_r_ 34,000 proinflammatory outer membrane protein (*oipA*) of *Helicobacter pylori*. Proceedings of the National Academy of Sciences, 97(13):7533–7538, 2000. doi: 10.1073/pnas.130079797.

Ziheng Yang. PAML 4: phylogenetic analysis by maximum likelihood. Molecular Biology and Evolution, 24(8):1586–1591, 2007. doi: 10.1093/molbev/msm088.

Ivan N. Zheludev, Robert C. Edgar, Maria Jose Lopez-Galiano, Marcos de la Peña, Artem Babaian, Ami S. Bhatt, and Andrew Z. Fire. Viroid-like colonists of human microbiomes. Cell, 187(23): 6521–6536.e18, 2024. doi: 10.1016/j.cell.2024.09.033.

